# The FDA-approved drug Nelfinavir inhibits lytic cell-free transmission of human adenoviruses

**DOI:** 10.1101/2020.05.15.098061

**Authors:** Fanny Georgi, Vardan Andriasyan, Robert Witte, Luca Murer, Silvio Hemmi, Lisa Yu, Melanie Grove, Nicole Meili, Fabien Kuttler, Artur Yakimovich, Gerardo Turcatti, Urs F Greber

## Abstract

Adenoviruses (AdVs) are prevalent and give rise to chronic and recurrent disease. The human AdV (HAdV) species B and C, such as HAdV-C2, C5 and B14, cause respiratory disease, and constitute a health threat for immuno-compromised individuals. HAdV-Cs are well known for lysing cells, owing to the E3 CR1-β-encoded adenovirus death protein (ADP). We previously reported a high-throughput image-based screening framework and identified an inhibitor of HAdV-C2 multi-round infection, Nelfinavir Mesylate. Nelfinavir is the active ingredient of Viracept, an FDA-approved inhibitor of the human immuno-deficiency virus (HIV) aspartyl protease, and used to treat acquired immunodeficiency syndrome (AIDS). It is not effective against single round HAdV infections. Here, we show that Nelfinavir inhibits the lytic cell-free transmission of HAdV, indicated by the suppression of comet-shaped infection foci in cell culture. Comet-shaped foci occur upon convection-based transmission of cell-free viral particles from an infected cell to neighbouring uninfected cells. HAdV lacking ADP was insensitive to Nelfinavir, but gave rise to comet-shaped foci indicating that ADP enhances but is not required for cell lysis. This was supported by the notion that HAdV-B14 and B14p1 lacking ADP were highly sensitive to Nelfinavir, although HAdV-A31, B3, B7, B11, B16, B21, D8, D30 or D37 were less sensitive. Conspicuously, Nelfinavir uncovered slow-growing round-shaped HAdV-C2 foci, independent of neutralizing antibodies in the medium, indicative of non-lytic cell-to-cell transmission. Our study demonstrates the repurposing potential of Nelfinavir with post-exposure efficacy against different HAdVs, and describes an alternative non-lytic cell-to-cell transmission mode of HAdV.

**Graphical Abstract:** Figure 1.

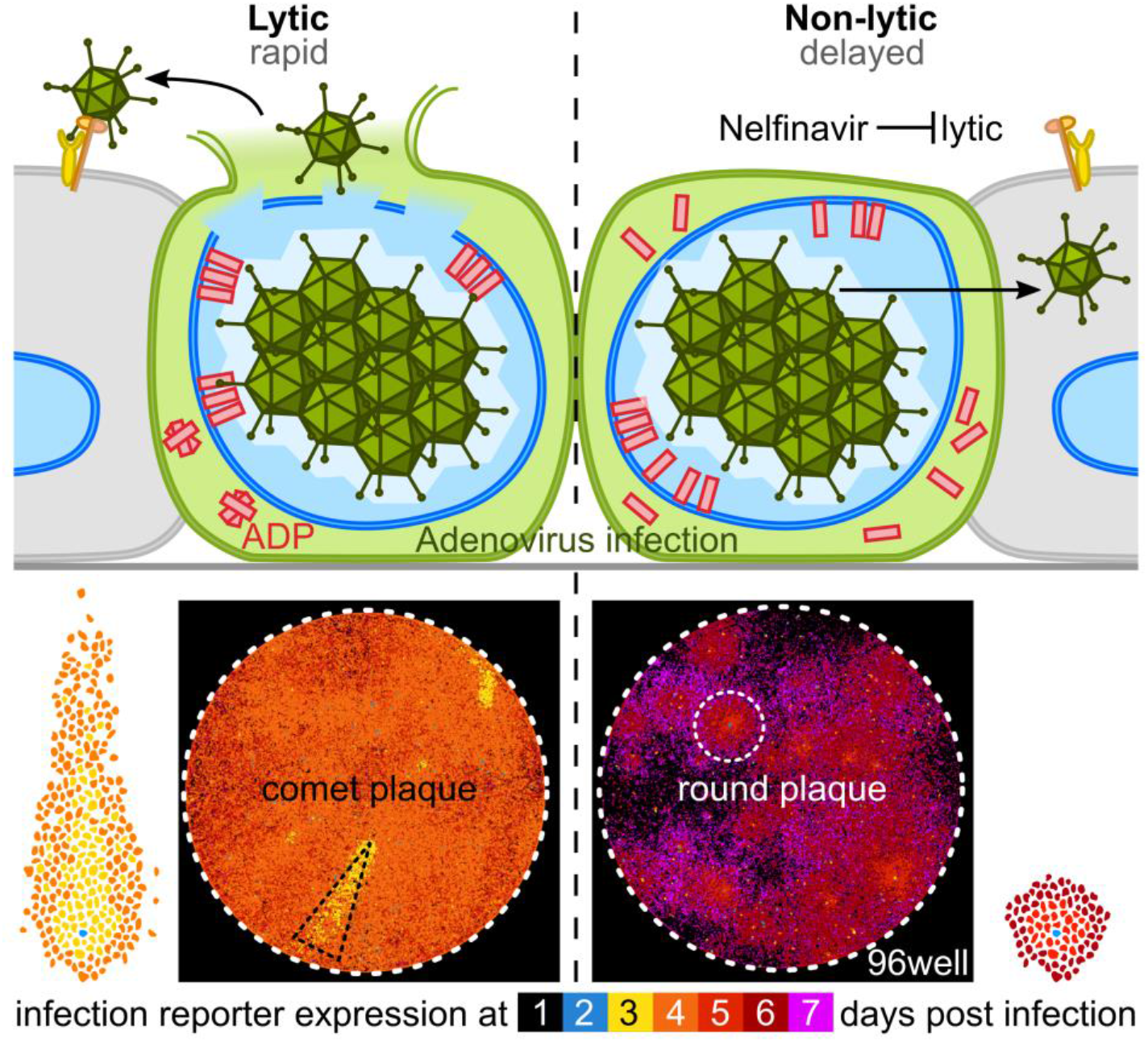

## Introduction

Adenovirus (AdV) was first described in 1953 by Rowe and co-workers as a cytopathologic agent isolated from human adenoids (Rowe et al., 1953). More than 100 human AdV (HAdV) genotypes have since been characterized by molecular genetics or serology and grouped into seven species (Harrach et al., 2019; Ismail et al., 2019). HAdV species A, F and G replicate in the gastrointestinal tract, B, C and E in the respiratory organs, and B and D in conjunctival cells of the eyes. Species B members have a broad tropism, including kidney and cells of the hematopoietic lineage (Lion, 2019; Lynch and Kajon, 2016; Mei et al., 2002). HAdV-caused illness can range from asymptomatic to lethal, especially in immunocompromised individuals (Bailey et al., 2018; Greber et al., 2013; Krilov, 2005). HAdV outbreaks are frequent in military training camps, but also nursery homes, as recorded in recurrent outbreaks of HAdV-E4 and HAdV-B7 (Erdman et al., 2002; Hwang et al., 2013; Lynch and Kajon, 2016; Potter et al., 2012; State of New Jersey Department of Health, 2019). To counter the disease burden, an oral HAdV-E4/B7 vaccine was reintroduced, leading to a sharp decline in adenoviral disease among military recruits (Deal et al., 2013; Lynch and Kajon, 2016; Radin et al., 2014). In addition to recurrent HAdV outbreaks, novel HAdV variants emerge, some of them causing pneumonia and death of elderly with chronic diseases. One of these emerging HAdVs is the HAdV-B14 variant 14p1, also known as 14a (Carr et al., 2011; Centers for Disease Control and Prevention (CDC), 2007; Lam et al., 2015; Louie et al., 2008; O’Flanagan et al., 2011). Furthermore, AdVs have a potential for zoonotic transmission (Benkő et al., 2014). Cross-species infections to humans from either non-human primates or psittacine birds have been reported from the USA and China, respectively (Chen et al., 2011b; To et al., 2014). Despite the high prevalence (Gray et al., 2007; Haque et al., 2018; Lynch and Kajon, 2016; Metzgar et al., 2007) and the broad use of AdV as gene therapy vectors (Ginn et al., 2018) as well as oncolytic viruses (Jiang et al., 2015; Lawler et al., 2017) no FDA-approved specific anti-HAdV treatment is available to date. Clinically, HAdV infections are treated with Ribavirin, Cidofovir, or more recently, Brincidofovir, which all inhibit viral DNA replication (Hiwarkar et al., 2017; Wold et al., 2019).

HAdV particles have been well characterized. They have a double-stranded DNA genome of ~36 kilo base pairs (kbp) packaged into an icosahedral capsid of about 90 nm in diameter (Benevento et al., 2014; Flatt and Greber, 2017; Liu et al., 2010; Reddy et al., 2010). HAdV-C2 and C5 replication cycle has been extensively studied including entry, uncoating, replication, assembly and egress from the infected cell (Allen and Byrnes, 2019; Atasheva et al., 2019; Charman et al., 2019; Greber, 2002; Greber and Flatt, 2019; Hidalgo et al., 2019; Kleinberger, 2019; Lynch et al., 2019; Nemerow and Flint, 2019; Oliveira and Bouvier, 2019; Pied and Wodrich, 2019; Prusinkiewicz and Mymryk, 2019; Sohn and Hearing, 2019; Suomalainen and Greber, 2013; Wang et al., 2018). HAdV-C infects cells by binding to the coxsackievirus adenovirus receptor (CAR) and integrin co-receptors, followed by receptor-mediated endocytosis, endosomal lysis and microtubule-motor driven transport to the nucleus, where it uncoats DNA and delivers the DNA into the nucleus (Bauer et al., 2019; Bremner et al., 2009; Burckhardt et al., 2011; Gastaldelli et al., 2008; Leopold et al., 2000; Luisoni et al., 2015; Meier and Greber, 2004; Suomalainen and Greber, 2013; Suomalainen et al., 1999; Wang et al., 2017; Wiethoff et al., 2005; Wolfrum and Greber, 2013; Zhou et al., 2018). The first viral protein expressed is E1A, a multifunctional intrinsically disordered protein controlling the transcriptional activity of all AdVs, as well as many cellular promoters, thereby affecting the cell cycle, differentiation, transformation and apoptosis (Berk, 2005; Ferrari et al., 2008; King et al., 2018; Pelka et al., 2008; Rao et al., 1992; Zemke and Berk, 2017). Viral early proteins besides E1A mediate immune escape, block activation of pro-apoptotic pathways and form nuclear viral DNA replication compartments. Late viral proteins give rise to mature progeny virions upon limited proteolysis of capsid proteins by the viral cysteine protease L3/p23 (Ahi and Mittal, 2016; Greber, 1998; Mangel and San Martín, 2014). Mature HAdV progeny is released upon rupture of the nuclear envelope and plasma membrane, which facilitates rapid viral dissemination and plaque formation *in vitro* (Doronin et al., 2003; Tollefson et al., 1996a; Yakimovich et al., 2012). The convection forces in the medium give rise to comet-shaped infection foci in cell cultures (Yakimovich et al., 2012). Foci of infected cells are also found in tissue, such as rat liver upon intravenous inoculation of HAdV-C5 (Haisma et al., 2008). Accordingly, acute HAdV infections trigger an inflammatory response, as shown in airways or conjunctiva of susceptible animals (Ismail et al., 2019; Kajon et al., 2003). In contrast to lytic virus transmission, direct cell-to-cell transmission leads to round plaques, as shown with vaccinia virus (Beerli et al., 2019; Doceul et al., 2010; Roberts and Smith, 2008; Zhong et al., 2013).

The mechanisms of virus transmission are highly virus-specific. They comprise non-lytic pathways involving the secretory-endocytic circuits, multi-vesicular or autophagic membrane processes, cellular protrusions, or transient breaches of membrane integrity (Burckhardt and Greber, 2009; Jansens et al., 2020; Mothes et al., 2010; van der Grein et al., 2018; Zhong et al., 2013). In contrast, lytic egress pathways further involve the destabilization of cellular membranes by viral and host factors, often tuned by the cytoskeleton (Danthi, 2016; Madan et al., 2008; Scott and Griffin, 2015; Wang et al., 2018; Zhang et al., 2018). HAdV-C2 controls lytic cell death by the adenovirus death protein (ADP), also known as 11.6K, as concluded from genetic and overexpression studies (Doronin et al., 2003; Tollefson et al., 1996a). ADP is a type III membrane protein transcribed from the CR1-β region in the immuno-regulatory E3a locus. All HAdV-C members harbour homologous E3a CR1-β sequences (e.g. 10.5K in HAdV-C5). Other HAdV species differ in their E3 region, however (Davison et al., 2003a; Dhingra et al., 2019; Robinson et al., 2013). The N-terminus of ADP is lumenal and the C-terminus protrudes into the cytosol (Scaria et al., 1992). Following post-translational modifications, ADP is transported to the inner nuclear membrane, where the N-terminus is intruding into the nucleus (Georgi and Greber, submitted). At late stages, when capsid assembly in the nucleus has commenced ADP expression is boosted (Tollefson et al., 1992; Wold et al., 1984). The mechanism of host cell lysis is still unknown, although necrosis-like, autophagic and caspase activities have been implicated (Abou El Hassan et al., 2004; Ito et al., 2006; Jiang et al., 2007, 2011).

Here, we report that Nelfinavir is an effective inhibitor of HAdV lytic egress. The identification process of Nelfinavir is described in an accompanying paper using an imaging-based, high content screen of the Prestwick Chemical Library (PCL) comprising 1,280 mostly clinical or preclinical compounds (Georgi et al., 2020; Yakimovich et al., 2015). Nelfinavir is the off-patent active pharmaceutical ingredient of Viracept, FDA-approved, which inhibits the human immuno-deficiency virus (HIV) protease (Kaldor et al., 1997). The work here documents the repurposing potential of Nelfinavir, which is effective against a spectrum of HAdV types in a post exposure manner. Nelfinavir is partly, but not exclusively, active against ADP-encoding HAdV types, and uncovers the appearance of round-shaped plaques, which arise upon non-lytic cell-to-cell viral transmission.

## Materials and Methods

### Viruses

HAdV-C2-dE3B-GFP was previously described (Yakimovich et al., 2012) (GenBank accession number MT277585). The virus was generated by exchange of the viral E3b genome region with a reporter cassette harbouring the enhanced green fluorescent protein (GFP) under a constitutively active cytomegalovirus (CMV) promoter. It was grown in A549 cells and purified by double CsCl gradient centrifugation (Greber et al., 1993). Aliquots supplemented with 10% (v / v) glycerol were stored at −80°C. HAdV-C2-dE3B-GFP was found to be homogeneous by SDS-PAGE and negative-stain analyses in transmission electron microscopy (EM). Recombinant HAdV-C2-dE3B-GFP-dADP was generated using homologous recombination according to the Warming recombineering protocols (Sirena et al., 2004; Warming et al., 2005). For a detailed protocol, see Supplementary methods. HAdV-C2-dE3B-GFP-dADP was plaque-purified and amplified, followed by two rounds of CsCl purification (Hemmi et al., 1998). Aliquots containing 10% (v / v) glycerol were stored at −80°C. HAdV-C2-dE3B-GFP-dADP was found to be homogeneous in SDS-PAGE and negative-stain analyses by transmission EM. Lack of ADP expression was confirmed by Western immunostaining using the rabbit α-HAdV-C2-ADP78-93 antibody, obtained from William Wold and Ann Tollefson (Saint-Louis University, Saint-Louis, USA) (Tollefson et al., 2003).

HAdV types A31, B7, B11, B14a, B16, B34, C1, C6, D8, D30 and D37 were kindly provided by the late Thomas Adrian (Hannover Medical School, Germany) and were verified by DNA restriction analysis (Adrian et al., 1986; Pacesa et al., 2017). HAdV types B14 (Carr et al., 2011; O’Flanagan et al., 2011) and B21a, isolate LRTI-6 (Kajon et al., 2015) were kindly provided by Albert Heim (Hannover Medical School, Germany). HAdV-B3-pIX-FS2A-GFP and B35-pIX-FS2A-GFP contain an enhanced GFP open reading frame (ORF) genetically fused to the downstream end of the HAdV pIX gene using an autocleavage FS2A sequence (Jetzer, 2018; Robinson et al., 2009; Studer, 2017). rec700 (Wold et al., 1986) and dl712 (Bhat and Wold, 1986) were obtained from William Wold (Saint-Louis University, Saint-Louis, USA). rec700 is a recombinant HAdV-C5 containing C2 sequences from nucleotide −236 to 2437 of the E3 transcription unit, and comprises the C2 E3a ORFs 12.5K, 6.7K, 19K and ADP, as well as major parts of the E3b ORF RID (10.4K protein) (Wold and Gooding, 1991). Mouse adenovirus (MAdV)-1-pIX-FS2A-GFP and MAdV-3-pIX-FS2A-GFP were constructed as described (Bieri, 2018; Hendrickx, 2016). HAdV-C2 and C5 were obtained from Maarit Suomalainen (University of Zurich, Switzerland). HSV-1-CMV-GFP is a recombinant HSV-1 strain SC16 containing a CMV enhancer/promoter-driven enhanced GFP expression cassette in the US5 (gJ) locus (Glauser et al., 2010) and was kindly provided by Cornel Fraefel (University of Zurich, Switzerland). HSV-1-CMV-GFP was propagated in Vero cells and purified by sucrose sedimentation as described in (Ali and Roossinck, 2007; Crameri et al., 2018). All viruses were stored in small aliquots containing 10% (v / v) glycerol at −80°C.

### Cell lines

A549 (human adenocarcinomic alveolar basal epithelium, CCL-185), HeLa (human epithelial cervix carcinoma, CCL-2) and HBEC (HBEC3-KT, normal human bronchial epithelium, CRL-4051) cells were obtained from the American Type Culture Collection (ATCC, Manassas, USA). HCE (normal human corneal epithelium) cells were obtained from Karl Matter (University College London, UK). CMT93 (mouse rectum carcinoma) cells were obtained from Susan Compton, Yale School of Medicine, USA. A549, HeLa, HCE and CMT-93 cell cultures were maintained in high glucose DMEM (Thermo Fisher Scientific, Waltham, USA) containing 7.5% (v / v) FCS (Invitrogen, Carlsbad, USA), 1% (v / v) L-glutamine (Sigma-Aldrich, St. Louis, USA) and 1% (v / v) penicillin streptomycin (Sigma-Aldrich, St. Louis, USA) and subcultured following phosphate-buffered saline (PBS) washing and trypsinisation (Trypsin-EDTA, Sigma-Aldrich, St. Louis, USA) bi-weekly. HBEC cells were maintained in endothelial-basal medium (ATCC, Manassas, USA) and passaged 1:1 weekly following PBS washing and trypsinisation. Cell cultures were grown at standard conditions (37°C, 5% CO_2_, 95% humidity) and passage number was limited to 20. Respective supplemented medium is referred to as supplemented medium.

### Compounds

Nelfinavir (CAS number 159989-65-8) powder was obtained from MedChemExpress LLC (Monmouth Junction, USA and Selleck Chemicals, Houston, USA). Compound was dissolved in DMSO (Sigma-Aldrich, St. Louis, USA) at 100 mM and kept at −80°C or −20°C for long-term or working storage, respectively.

### Cellular impedance measurement

Impedance-based assays were performed using the xCELLigence system (Roche Applied Science and ACEA Biosciences) as described previously (Prasad et al., 2014, 2020a) according to the manufacturer’s instructions (Spiegel, 2009) in cell culture environment (37°C, 5% CO_2_, 95% humidity) in duplicates. The 16-well E plates have a gold-plated sensor array embedded in their glass bottom by which the electrical impedance across each well bottom is measured. The impedance per well termed cell index (CI) is recorded as a dimensionless quantity. The background CI was assessed following the addition of 50 μl supplemented medium to each well and equilibration in the incubation environment. After 30 min equilibration, 9,000 A549 ATCC cells in 50 μl supplemented medium were added per well and measurement was started.

For the quantification of Nelfinavir toxicity, 50 μl supernatant was removed 18 h later and replaced by 2-fold concentrated Nelfinavir or DMSO solvent as the control dilution in supplemented medium (final Nelfinavir concentration 0.4-100 μM in 100 μl / well). The control was supplemented medium. Impedance was recorded every 15 min over 5 days. Cytotoxicity of Nelfinavir over time is given as toxic concentration 50%, TC_50_. It indicates the concentration at which the cumulated impedance of the treated cells is twice as high as background impedance levels. TC_50_ was calculated by non-linear regression of solvent-normalized CI over the concentration of Nelfinavir.

For the quantification of Nelfinavir effects on the cytopathogenicity of HAdV-C2-dE3B-GFP compared to HAdV-C2-dE3B-GFP-dADP infection, 50 μl supernatant were removed 18 h later and replaced with Nelfinavir- and virus-supplemented medium. 25 μl of a 4-fold concentrated Nelfinavir (final concentration 0.4-100 μM) or corresponding DMSO solvent control dilution (final concentration 1%) in supplemented medium or supplemented medium only were added to 50 μl medium containing cells. Additionally, 25 μl of a 4-fold concentrated virus stock dilution were added (final inoculum 1.68*106 viral particle(s) (VP) / well HAdV-C2-dE3B-GFP and 2.68*106 VP/ well HAdV-C2-dE3B-dADP, corresponding to ~30 plaque forming unit(s) (pfu) / well). The delay of infection-induced cytotoxicity was calculated as time point at which the CI of the infected cells had decreased by 50% relative to its maximum. Data analysis was performed using GraphPad (GraphPad Software, Inc, version 8.1.2), and curve fitting was performed using three-parameter [inhibitor] vs. response nonlinear regression.

### Fluorescence-based plaque forming assay

Per 96-well, 15,000 A549, 10,000 HeLa ATCC, 30,000 HBEC, 30,000 HCE or 30,000 CMT-93 cells were seeded in 100 μl of the respective supplemented medium and allowed to settle for 1 h at room temperature (RT) prior to cell culture incubation at 37°C, 5% CO_2_, 95% humidity. The following day, the medium was replaced by 50 μl of the respective virus stock dilution giving rise to 5 to 50 plaques per 96-well. 50 μl Nelfinavir to obtain 0.1 to 50 μM final concentration or DMSO solvent control was also added, both in supplemented medium. For each experiment, a non-infected, treated control was performed. For uphill plaque assays, medium volume was increased to 150 μl with identical virus and drug concentrations. For wash-in/wash-out experiments, virus was incubated on the cells in supplemented medium for 1 h at 37°C, cells were washed with PBS and 100 μl drug dilution in supplemented medium was added. All experiments were performed in four technical replicates or as indicated. Cells were incubated at standard cell culture conditions. At the indicated time post infection (pi), the cells were fixed and the nuclei stained for 1 h at RT by addition of 33 μl 16% (w / v) para-formaldehyde (PFA)and 4 μg / ml Hoechst 33342 (Sigma-Aldrich, St. Louis, USA) in PBS. Cells were washed three times with PBS and stored in PBS supplemented with 0.02% N3 for infections with viruses harbouring a GFP transgene. For wild type (wt) viruses, cells were quenched in PBS supplemented with 50 mM NH_4_Cl, permeabilized using 0.2% (v / v) Triton-X100 in PBS and blocked with 0.5% (w / v) BSA in PBS. Cells were incubated with 381.7 ng / ml mouse α-HAdV hexon protein antibody (Mab8052, Sigma-Aldrich, St. Louis, USA) and subsequently stained using 2 μg / ml goat α-mouse-AlexaFluor594 (A21203 or A32742, Thermo Fisher Scientific, Waltham, USA). Plates were imaged on either an IXM-XL or IXM-C automated high-throughput fluorescence microscope (Molecular Devices, San Jose, USA) using a 4x objective at widefield mode. Hoechst staining was recorded in DAPI channel, FITC / GFP channel for viral GFP and TRITC / Texas red channel for immunofluorescence hexon staining.

### Therapeutic index measurement

The infection phenotype for each well was quantified using Plaque2.0 (Yakimovich et al., 2015). The number of plaques was determined based on the infection signal (viral GFP or hexon immunofluorescence staining). Nuclei were segmented based on Hoechst signal by CellProfiler (Carpenter et al., 2006). Hereof infected nuclei were classified based on the median infection signal per nucleus in CellProfiler. Data were plotted and EC50 (infected and treated cells), TC_50_ (non-infected, treated cells), as well as the corresponding standard error (SE) determined based on curve fitting in GraphPad (GraphPad Software, Inc, version 8.1.2) using three-parameter [inhibitor] vs. response nonlinear regression. Mean TI_50_ was calculated as EC50 / TC_50_ ratio of the means. The TI_50_ SE is calculated by error propagation.

### Quantification of viral protein expression

Infection, HAdV hexon immunofluorescence staining, and imaging were performed as described under Microscopic plaque assay in technical quadruplicates. Single nuclei were segmented based on the Hoechst signal, using CellProfiler (Carpenter et al., 2006). Median GFP and hexon signal per nucleus were measured and infected nuclei were classified based on the median GFP signal per nucleus and. Subsequently, mean and standard deviation (SD) over all infected nuclei per well were calculated in R version 3.3.2 (R Core Team, 2018). Data were plotted in GraphPad (GraphPad Software, Inc, version 8.1.2).

### Transmission electron microscopy

A549 ATCC cells grown on Alcian Blue-treated cover slips were infected with HAdV-C2-dE3B-GFP in supplemented medium with 0, 1.25 or 3 μM Nelfinavir and cultured for 40 h at standard cell culture conditions. The samples were washed with ice-cold 0.1 M cacodylate buffer (pH 7.4) and fixed at 4°C in 0.1 M ice-cold cacodylate buffer (pH 7.4), supplemented with 2.5% (v / v) glutaraldehyde and 0.5 mg / ml ruthenium red for 1 h. Cells were washed with 0.1 M cacodylate buffer (pH 7.4) and post-fixed at RT in 0.05 M cacodylate buffer (pH 7.4) supplemented with 0.5% (v / v) OsO_4_ and 0.25 mg / ml ruthenium red for 1 h. Following washing with 0.1 M cacodylate buffer (pH 7.36) and H_2_O, the samples were incubated in 2% (v / v) uranyl acetate at 4°C over night (ON). The samples were dehydrated in acetone and embedded in Epon as described in (Greber et al., 1997). 85 nm slices were obtained (Leica Ultracut UCT, Leica, Wetzlar, Germany) and stained with uranyl acetate.

### HAdV-C5 virus production in presence of Nelfinavir

HAdV-C5 was amplified in the medium containing 0, 1.25 or 3 μM Nelfinavir for 4 days. Cells were harvested and disrupted by three freeze / thaw cycles. The cell debris was removed by Freon extraction and mature full HAdV virions were purified by two rounds of CsCl gradient ultracentrifugation (Hemmi et al., 1998). Protein concentration was determined by BCA assay (Pierce BCA Protein Assay Kit, Thermo Fisher Scientific, Waltham, USA). For long-term storage, virus stocks were supplemented with 10% (v / v) glycerol and kept at −80°C.

### Negative staining electron microscopy

Double CsCl gradient-purified HAdV particles were adhered to Collodion and 2% (v / v) amyl acetate film-covered grids (300 mesh Formvar/carbon-supported copper support films, Electron Microscopy Sciences, Hatfield, USA). Viral particles were negatively stained with 2% (v / v) uranyl acetate and viewed on a transmission electron microscope (Philips CM100, Philips, Amsterdam, Netherlands) at 100 kV. Images were acquired using a CCD camera (Orius SC1000 with 4,000 x 2,600 pixels, Gatan, Pleasanton, USA).

### Western blot analysis of HAdV protease activity

Double CsCl-purified grown in presence / absence of Nelfinavir (HAdV-C5^±Nelfinavir^) stocks and size standard (PageRuler plus, Thermo Fisher Scientific, Waltham, USA) were size-separated on 12% acrylamide gel under reducing conditions and transferred to a PVDF membrane. HAdV proteins were detected using the following primary antibodies: 1:10,000 R72 rabbit α-fiber (Baum et al., 1972), 1:1,000 rabbit α-pVI/VI (Burckhardt et al., 2011), 1:1,000 R3 rabbit α-pVII/VII (Ulf Pettersson of Uppsala University) and visualized using a goat α-rabbit-HRP (7074, Cell Signaling Technology, Danvers, USA) and ECL Prime Western Blotting Detection Reagent (GE Health Care, Pittsburgh, USA). The membranes were luminescence imaged on an Amersham Imager 680 (GE Health Care, Pittsburgh, USA).

### Determination of nuclear size

Infection and Nelfinavir treatment of A549 cells were performed as described under Microscopic plaque assay with a cell seeding density of 15,000 cells / well. Wells were imaged with IXM-C automated high-throughput fluorescence microscope (Molecular Devices, San Jose, USA) using a 40x objective (NA 0.95) at confocal mode (62 μm pinhole). DAPI channel was acquired for nuclear Hoechst staining, FITC / GFP channel was acquired for viral GFP, TRITC / Texas red channel was acquired for immunofluorescence ADP staining and Cy5 channel was acquired for NHS-ester signal. 30 z steps with 0.5 μm step size were acquired for each channel and maximal projections were calculated. Image analysis was performed using CellProfiler (Carpenter et al., 2006). Nuclei areas were segmented based on thresholded Hoechst signal. Infected cells were classified based on a fixed threshold for median nuclear GFP intensity. Data processing was performed in R version 3.3.2 (R Core Team, 2018). Statistical analysis was performed in GraphPad (GraphPad Software, Inc, version 8.1.2) using the non-parametric Kolmogorov-Smirnov test.

### Cell binding assay of virus

A549 cells were seeded at 7,500 cells per 96-well in full DMEM and allowed to attach over night at standard cell culture conditions. The next day, the medium was replaced by 3*108 VP/ well of double CsCl-purified HAdV-C5^±Nelfinavir^ virus stocks in 100 μl ice-cold supplemented medium and kept on ice for 30 min. Following a 15 min entry phase under standard cell culture conditions the cells were fixed and the nuclei stained for 1 h at RT by addition of 33 μl 16% PFA and 4 μg / ml Hoechst 33342 (Sigma-Aldrich, St. Louis, USA) in PBS. Following the above described immuno-fluorescence staining procedure, the cell-bound HAdV virions were stained using 9C12 mouse α-hexon (developed by Laurence Fayadat and Wiebe Olijve, obtained from Developmental Studies Hybridoma Bank developed under the auspices of the National Institute of Child Health and Human Development and maintained by the University of Iowa, Iowa City, USA) (Varghese et al., 2004) and goat α-mouse AlexaFluor488 (A11029, Thermo Fisher Scientific, Waltham, USA). Total area was identified by Alexa-Fluor647 NHS ester staining (A20006, Thermo Fisher Scientific, Waltham, USA). Max projections of confocal z-stacks (25 z steps spaced 1 μm) were acquired on a SP5 resonant APD (Leica, Wetzlar, Germany) at 1.7x zoom using a 63x glycerol objective (numerical aperture 1.4).

### Assessment of HAdV infectivity of HAdV-C5^±Nelfinavir^

Fifteen thousand A549 cells were seeded per 96-well in full DMEM and allowed to attach over night at standard cell culture conditions. The next day, the medium was replaced by double CsCl-purified HAdV-C5^±Nelfinavir^ virus stocks at 50 to 0.001 pg / well of BCA-based viral protein concentration and incubated at standard cell culture conditions. Cells were fixed at 52 hpi, stained for HAdV hexon expression and imaged following the procedure described under Image-based plaque assay. Images were quantified using Plaque2.0 (Yakimovich et al., 2015). Nuclei were segmented based on Hoechst signal. Infected cells were segmented based on hexon immunofluorescence staining signal.

### Egress assay

A549 cells were seeded at 480,000 cells per 6-well in full DMEM and infected at 1,100 pfu HAdV-C2-dE3B-GFP per well the next day. Following 1 h of warm incubation, the supernatant was removed, and cells were washed with PBS and detached by trypsin digestion. Infected cells were centrifuged and resuspended in fresh medium to remove any unbound input virus and seeded at 180,000 cells / 12-well in medium supplemented with 1.25, 3 or 10 μM Nelfinavir or equivalent amounts of DMSO solvent control. At the indicated times pi, the supernatant was harvested and cleared by centrifugation. 200 μl PBS / well was added to the infected monolayer. Cells were disrupted by three freeze / thaw cycles and freon extraction was performed. Supernatant and cell lysate were stored at 4°C until titration on naive A549 cells. PFA-fixed, Hoechst-stained cells were imaged at 44 hpi using a 4x objective (NA 0.20) on an epifluorescence IXM-XL (Molecular Devices, San Jose, USA). GFP-positive infected cells were classified based on median nuclear GFP intensity using automated image analysis by CellProfiler (Carpenter et al., 2006).

### Quantification of infectious progeny production

Four hundred and eighty thousand A549 cells were seeded per 6-well dish and inoculated with 1,100 pfu HAdV-C2-dE3B-GFP / well for 1 h at 37°C, washed with PBS and detached by trypsin digestion. Infected cells were centrifuged and resuspended in fresh medium to remove any unbound input virus. Cells were seeded at 180,000 cells / 12-well in medium supplemented with 1.25, 3 or 10 μM Nelfinavir or the respective DMSO solvent control. Viral progeny in the cell monolayer and supernatant was harvested at the indicated time pi by three freeze / thaw cycles. The lysates were cleared by centrifugation and stored at 4°C until titration on naive A549 cells. PFA-fixed, Hoechst-stained cells were imaged at 44 hpi using a 4x objective on an epifluorescence IXM-XL (Molecular Devices, San Jose, USA). GFP-positive infected cells were classified based on median nuclear GFP intensity using automated image analysis by CellProfiler (Carpenter et al., 2006). The yield per 12-well was extrapolated by linear regression of the number of infected cells per μl of harvested whole well lysate using GraphPad (GraphPad Software, Inc, version 8.1.2).

### Quantification of the antiviral potency of Nelfinavir

Infection was performed as described under Microscopic plaque assay. Cells were incubated with an inoculum ranging between 10 - 2,560 pfu / well HAdV-C2-dE3B-GFP for 1 h at 37°C. Cells were washed with PBS and 100 μl DMEM phenol-free medium (Thermo Fisher Scientific, Waltham, USA), supplemented with 1% penicillin streptomycin (Sigma-Aldrich, St. Louis, USA), 1% L-glutamine (Sigma-Aldrich, St. Louis, USA), 7.5% FBS (Invitrogen, Carlsbad USA), 1% non-essential amino acids (Sigma-Aldrich, St. Louis, USA), 1% 100 mM sodium pyruvate (Thermo Fisher Scientific, Waltham, USA), 0.25 ng / ml Hoechst 33342 (Sigma-Aldrich, St. Louis, USA) and 1 μg / ml propidium iodide (PI, Molecular Probes, Eugene, USA). Plates were imaged at the indicated times pi on an IXM-C automated high-throughput fluorescence microscope (Molecular Devices, San Jose, USA) using a 40x objective (NA 0.95) at confocal mode (62 μm pinhole). DAPI channel was acquired for nuclear Hoechst staining, FITC / GFP channel was acquired for viral GFP and Cy5 channel was acquired for PI signal. 30 z steps with 0.5 μm step size were acquired for each channel and maximal projections were calculated.

### Morphological plaque characterization

Plaques were segmented in Plaque2.0 (Yakimovich et al., 2015) and plaque region eccentricity was measured as fraction of the distance between the two focal points of the ellipse divided by the length of the major axis. Only plaque regions consisting of at least five infected cells (≥6,000 px2) with a centroid located 600 px from the well rim were considered to exclude spatial limitations. Plaque roundness was calculated as 1-eccentricity (Equation 1).

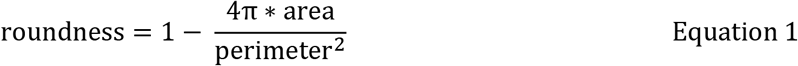

Statistical analysis was performed in GraphPad (GraphPad Software, Inc, version 8.1.2) using the non-parametric Kolmogorov-Smirnov test.

### Confocal microscopy of ADP localization

Infection and immunofluorescence stainings were performed as described under Microscopic plaque assay with a cell seeding density of 3,000 cells / well. Cells were incubated with 1:1,000 rabbit α-HAdV-C2-ADP87-101 antibody (Tollefson et al., 2003) and subsequently stained using donkey α-rabbit-AlexaFluor594 (21207, Thermo Fisher Scientific, Waltham, USA) and 0.2 μg/ml NHS ester (Life Technologies, Carlsbad, USA) for whole cell outline. Plates were imaged on an IXM-C automated high-throughput fluorescence microscope (Molecular Devices, San Jose, USA) using a 40x objective (NA 0.95) at confocal mode (62 μm pinhole). DAPI channel was acquired for nuclear Hoechst staining, FITC / GFP channel was acquired for viral GFP, TRITC / Texas red channel was acquired for immunofluorescence ADP staining and Cy5 channel was acquired for NHS ester signal. 30 z steps with 0.5 μm step size were acquired for each channel and maximal projections were calculated. Image analysis was performed using CellProfiler (Carpenter et al., 2006). Nuclei and whole cell areas were segmented based on thresholded Hoechst and NHS ester signal, respectively. Nuclear rim was defined as 10 pixel-wide area around the nuclear border. Infected cells were classified based on the whole cell 5% quantile GFP intensity. Whole cell and nuclear rim mean TRITC / Texas red (detecting ADP) intensities as well as whole cell 5-pixel granularity per infected cell were normalized by the according mean over all infected cells of the solvent control. Data processing was performed in R version 3.3.2 (R Core Team, 2018). Statistical analysis was performed in GraphPad (GraphPad Software, Inc, version 8.1.2) using the non-parametric Kolmogorov-Smirnov test.

### Western blot analysis of ADP processing

Four hundred and eighty thousand A549 cells were seeded per 6-well, incubated o/n and inoculated with HAdV-C2-dE3B-GFP at 22,000 pfu / well in 1.2 ml full DMEM supplemented with 0 to 10 μM Nelfinavir. Following 44 h of incubation in standard cell culture medium, cells were placed on ice and the supernatant was removed. The cells were washed twice with ice-cold PBS. Cells were lysed in 100 μl COS lysis buffer (20 mM Tris-HCl pH 7.4, 100 mM NaCl, 1 mM EDTA, 1% Triton X-100, 1 mM DTT, 25 mM β-Glycerophosphate disodium, 25 mM NaF, 1 mM Na3VO4, 1x protease inhibitors (Mini Complete, Roche, Basel, Switzerland) for 5 min on ice. Supernatant and washing PBS were collected and cells pelleted by centrifugation at 16,000 xg for 5 min at 4°C. Lysates were scraped off and used to resuspend the pelleted cells. Following another centrifugation, the supernatant was collected and stored at −20°C. Samples of 15 μl lysate were supplemented with SDS-containing loading buffer (0.35 M Tris-HCl pH 6.8, 0.28% SDS, 30 g / l DTT, 0.6 g / l bromophenol blue). Samples were denatured at 95°C for 5 min and proteins were separated on a denaturing 15% acrylamide gel. Proteins transferred to a PVDF membrane were detected with 1:1,000 of a rabbit α-HAdV-C2 ADP78-93 antibody (Tollefson et al., 2003) followed by goat α-rabbit-HRP (7074, Cell Signaling Technology, Danvers, USA). Protein bands were visualized using ECL Prime Western Blotting Detection Reagent (GE Health Care, Pittsburgh, USA) and luminescence imaged on an Amersham Imager 680 (GE Health Care, Pittsburgh, USA).

### Neutralization of HAdV cell-free progeny

A549 cells were seeded at 15,000 cells per well of a 96-well-plate, incubated o/n and inoculated with HAdV-C2-dE3B-GFP at 34 pfu / well for 1 h at 37°C. Virus was removed and cells were washed with PBS, before 0.25 ng / ml Hoechst (Sigma-Aldrich, St. Louis, USA)-supplemented DMEM medium containing 1:12 HAdV-C2/5-neutralizing dog serum, kindly supplied by Anja Ehrhardt, University Witten/Herdecke, Germany (Hausl et al., 2010), supplemented with 40% v / v glycerol), control goat serum (Thermo Fisher Scientific, Waltham, USA, supplemented with 40% v / v glycerol) or the corresponding volume glycerol only. Cells were imaged using a 4x objective (NA 0.20) on an epifluorescence IXM-XL microscope (Molecular Devices, San Jose, USA).

### Crystal violet-stained plaques

Plaque shapes were also assessed by conventional crystal violet-stained plaque assay, performed in A549 cells in liquid supplemented DMEM medium. All infections were performed at 37°C, 95% humidity and 5% CO_2_ atmosphere. At the indicated time pi, cells were fixed and stained for 60 min with PBS solution containing 3 mg / ml crystal violet and 4% PFA added directly to the medium from a 16% stock solution. Plates were de-stained in H_2_O, dried and imaged using a standard 20 mega pixel phone camera under white light illumination.

## Results

### Nelfinavir is a non-toxic, potent inhibitor of HAdV-C multicycle infection

An accompanying paper describes a full cycle, image-based screen of 1,278 out of 1,280 PCL compounds against HAdV-C2-dE3B-GFP, where Clopamide and Amphotericine B were excluded due to precipitation during acoustic dispension into the screening plates (Georgi et al., 2020). The screen was conducted in adenocarcinomic human alveolar basal epithelial (A549) cells at 1.25 μM compound concentration, and identified Nelfinavir, Aminacrine, Dequalinium dichloride and Thonzonium bromide as hits (Supplementary Table 1). Nelfinavir (CAS number 159989-65-8), hereafter referred to as Nelfinavir, strongly inhibited plaque numbers at nanomolar concentrations, comparable to the known HAdV nucleoside analogue inhibitor 3’-deoxy-3’-fluorothymidine (DFT, Figure 1A, 1B). Dequalinium dichloride, Aminacrine and Thonzonium bromide were excluded from further analyses due to toxicity (Georgi et al., 2020), and potential mutagenic effects (Topal, 1984). Long-term incubations of uninfected A549 cells with Nelfinavir up to 115 h showed median toxicity TC_50_ of 25.7 μM, as determined by cell impedance measurements using xCELLigence (Figure 1C), consistent with presto-blue assays and counts of cell nuclei (Supplementary Figure 1A, Supplementary Table 1). This was in agreement with previous reports, and acceptable side effects in clinical use against HIV (Kaldor et al., 1997; Moyle et al., 1998). The therapeutic index 50 (TI_50_) of Nelfinavir was 27.1 (Figure 1D), as determined by the ratio between the concentration yielding 50% loss of cell nuclei (TC_50_ = 10.01 μM) and the effective concentration yielding 50% inhibition of fluorescent plaque formation (EC_50_ = 0.37 μM). The data indicates that Nelfinavir is an effective, non-toxic inhibitor of HAdV-C2 multi-cycle infection.

**Figure 1.**
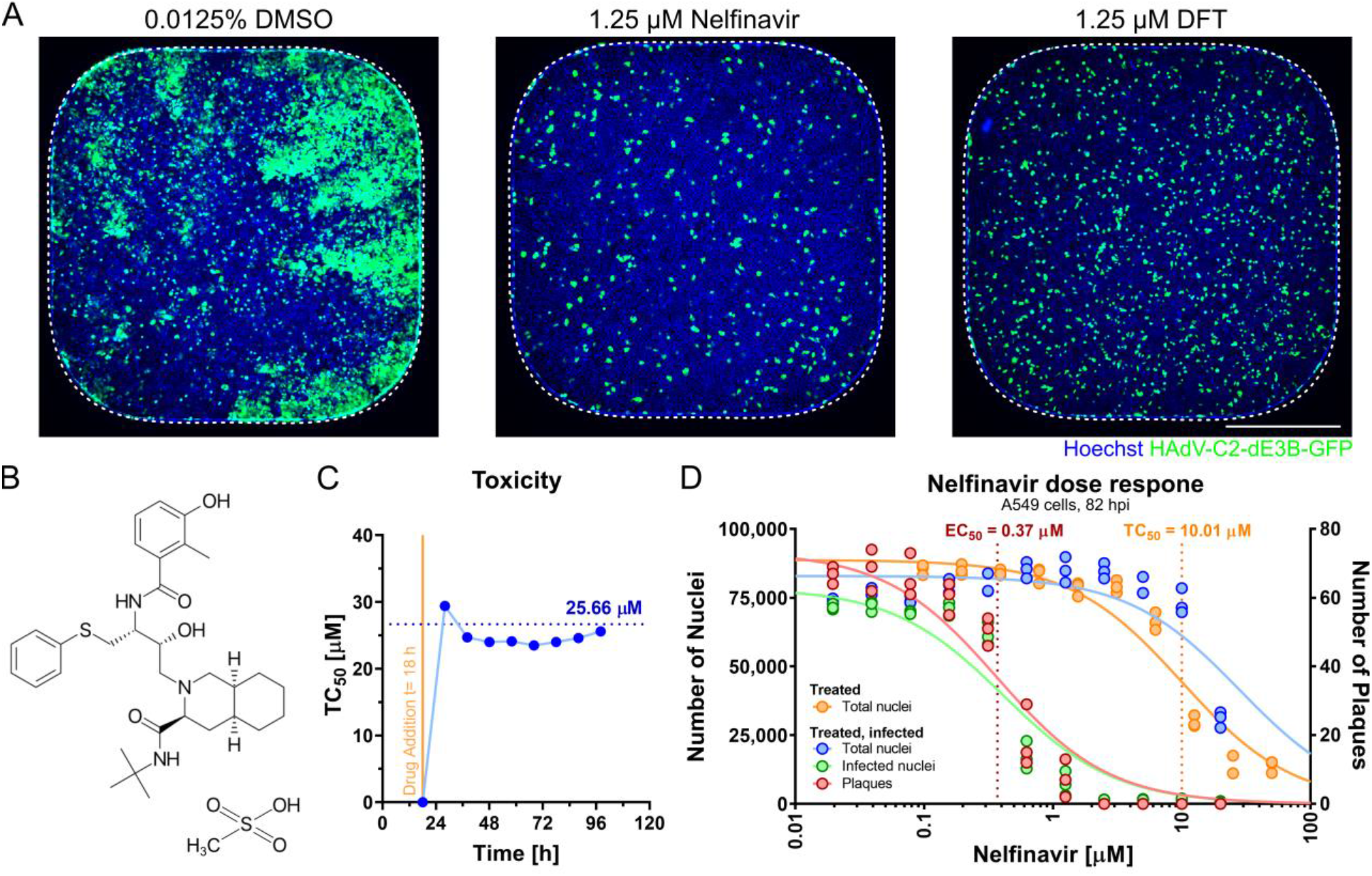
The small molecule Nelfinavir is a potent inhibitor of HAdV-C infection. **A** Representative 384-well epifluorescence microscopy images of cells treated with DMSO (left), Nelfinavir (centre) and DFT (right), and infection with HAdV-C2-dE3B-GFP for 72 h. Hoechst-stained nuclei are shown in blue, viral GFP in green. Dotted lines indicate well outline. Scale bar is 5 mm. **B** Structural formula of Nelfinavir Mesylate. **C** Cell index (CI)-based concentration causing 50% toxicity in uninfected cells (TC_50_) upon long-term incubation of A549 with Nelfinavir. Impedance was recorded every 15 min using an xCELLigence system. The time on the x-axis indicates hours after cell seeding. Vertical line shows the time of drug addition (raw data available in Supplementary Figure 1). **D** Separation of effect (EC_50_, plaque numbers) and toxicity (TC_50_, nuclei numbers) of Nelfinavir in A549 cells at 82 hpi based on four technical replicates. Plaque numbers per well are depicted as red circles, and numbers of infected nuclei as green circles. Numbers of nuclei in Nelfinavir-treated, uninfected wells are shown in blue; treated, infected wells shown in orange.

### Nelfinavir does not affect single round infection

We first tested if Nelfinavir affected viral protein production. HAdV-C2-dE3B-GFP-infected A549 cells were analysed for GFP under the immediate early CMV promoter, and the late protein hexon expressed after viral DNA replication at 46 hours post infection (hpi). Results indicate that Nelfinavir had no effect on GFP or hexon expression at the tested concentrations, while the formation of fluorescent plaques was completely inhibited (Figure 2A, and Figure 1D). This result was in agreement with the notion that Nelfinavir did not affect the replication of the HAdV-C5 genome, as determined by q-PCR (Gantt et al., 2011). We next examined if Nelfinavir affected the formation of viral particles. Transmission electron microscopy (TEM) of HAdV-C2-dE3B-GFP-infected cells revealed large numbers of virions in the nuclei of Nelfinavir-treated and untreated cells (Figure 2B). This result was conforming with the observation that the nuclei of Nelfinavir-treated cells expanded in area over time, indistinguishable from control cells (Supplementary Figure 1B).

**Figure 2.**
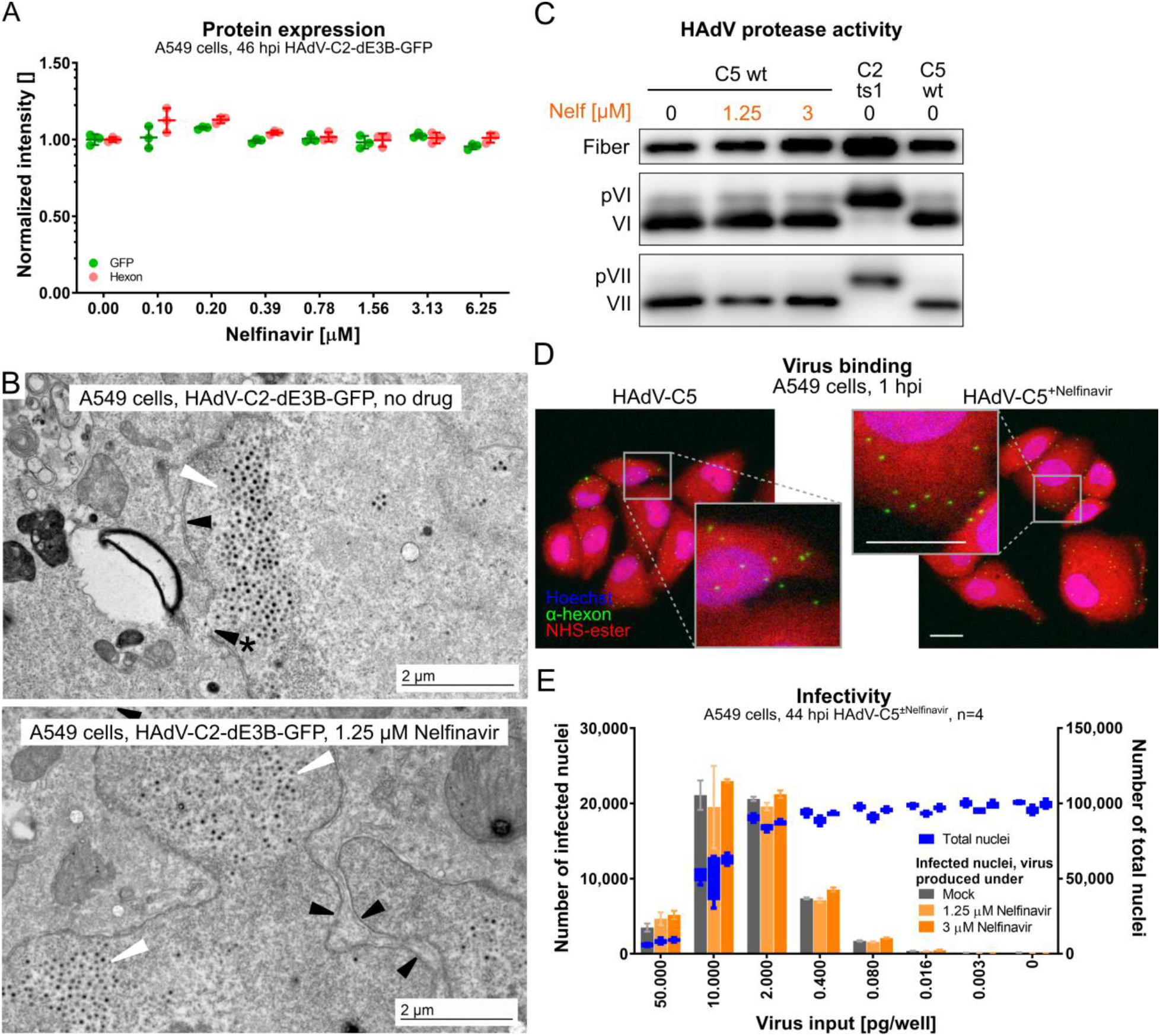
Nelfinavir does not affect early or late steps of HAdV-C infection. **A** No effects of Nelfinavir on the expression level of CMV-GFP (green) or the late HAdV protein hexon (red) in HAdV-C2-dE3B-GFP-infected A549 cells. Data points represent for each of the four biological replicates: mean median, nuclear intensities per well normalized to the mean median nuclear intensities of the DMSO-treated wells. Epifluorescence microscopy images were segmented and analysed using CellProfiler. **B** Representative TEM images of late stage HAdV-C2-dE3B-GFP-infected A549 cells at 41 hpi reveal viral particles inside the nucleus in both DMSO-treated and Nelfinavir-treated cells (white arrow head). Black arrow heads indicate the nuclear envelope, arrow head with * points to rupture. Scale bar equivalent to 2 μm. **C** Nelfinavir does not affect the maturation of HAdV-C5, as indicated by fully processed VI and VII proteins in purified particles grown in presence of Nelfinavir. Note that HAdV-C2-ts1 lacking the L3/p23 protease contains the precursor capsid proteins of VI and VII (pVI and pVII). **D** HAdV-C5 grown in presence of Nelfinavir (HAdV-C5^+Nelfinavir^) binds to naive A549 cells similar as HAdV-C5 from control cells. Cells were incubated with virus at 4°C for 1 h and fixed with PFA. Staining with Hoechst (blue nuclei) and for virus capsids with an α-hexon antibody (green puncta). Cell material was visualized by NHS-ester staining (red signal). Images are max projections of confocal z stacks, and also show zoomed in views (grey squares). Scale bars equivalent to 20 μm. **E** Particles produced in presence of Nelfinavir are fully infectious. A549 cells were inoculated with purified HAdV-C5 grown in presence or absence of Nelfinavir. Number of infected cells at 44 hpi, shown in grey and orange, were quantified based on α-hexon immunofluorescence staining. Total cell numbers were segmented based on nuclear Hoechst staining (blue). Bars represent means of four technical replicates. Error bars indicate standard deviation.

To test if Nelfinavir affected virion maturation, we analysed purified virions by SDS-PAGE and Western blotting against proteins pVI/VI and pVII/VII using previously characterized antibodies. There was no evidence for increase of precursor VI or VII (pVI or pVII) in HAdV-C5 from Nelfinavir-treated cells, in contrast to temperature-sensitive (ts) 1 particles, which lack the L3/p23 protease due to the point mutation P137L in p23 (Imelli et al., 2009) (Figure 2C). This showed that Nelfinavir did not affect the proteolytic maturation of the virus by the L3/p23 cysteine protease. In accordance, purified HAdV-C5 from Nelfinavir-treated cells attached to naive A549 cells and gave rise to viral gene expression as effectively as control HAdV-C5 particles (Figures 2D, 2E). Together, these results indicate that Nelfinavir does not affect the production of infectious virions in single round infections.

### Nelfinavir inhibits HAdV-C egress

We investigated the kinetics of HAdV-C2-dE3B-GFP production and the release to the supernatant. Supernatants and whole cell lysates of treated- and non-treated infected cells were harvested at different time points. Unperturbed cells started to lyse 44 hpi, second-round infections were well advanced to plaques at 72 hpi, and most of the infected cells had lysed and released progeny at 120 hpi (Figure 3A). At 44 hpi, cell lysates of Nelfinavir- and control-treated cells contained similar infectivity, and supernatants were essentially free of virus, as shown by titration on naive A549 cells. At 72 hpi, supernatants of Nelfinavir-treated cells contained no infectious virus, while the supernatant of non-treated cells contained infectious particles. Similar findings were made at 120 hpi comparing the supernatant of cells treated with 3 μM Nelfinavir to non-treated cells. The difference in infectious load was confirmed by titration of supernatants from separate time course experiments at three different concentrations of Nelfinavir (Figure 3B). At 7 dpi, a dosage of 1.25 μM reduced the total yield of infectious particles in the supernatant by three orders of magnitude, underscoring the potency of Nelfinavir to block the dissemination of HAdV-C-dE3B-GFP. Moreover, Nelfinavir limited HAdV-C2 transmission when added as late as 40 hpi (Figure 3C). These findings indicate that Nelfinavir impairs the egress of progeny from the host cell.

**Figure 3.**
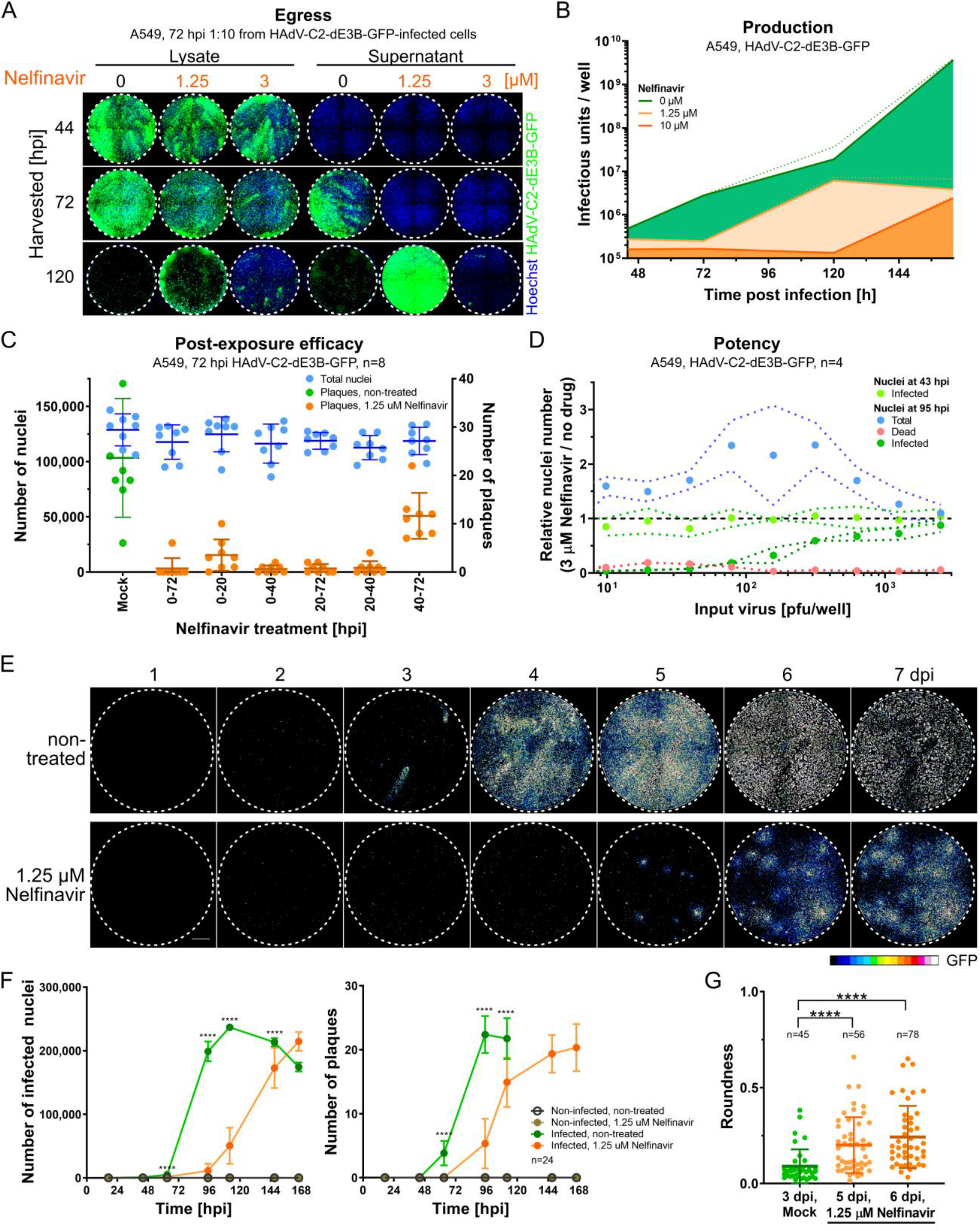
Nelfinavir is a post-exposure inhibitor of HAdV-C egress. **A** A549 indicator cells were inoculated with 1:10 diluted cell lysates or supernatants from Nelfinavir or control-treated A549 cells, which had been infected with HAdV-C2-dE3B-GFP for indicated durations, and incubated for 3 days. Results reveal delayed viral progeny release to the supernatant of Nelfinavir-treated cells. Nuclei signal shown in blue, viral GFP in green. **B** Released and cell-associated total progeny from HAdV-C2-dE3B-GFP-infected A549 cells treated with Nelfinavir (orange) or DMSO (green) determined by titration on naive A549 cells in a 12-well assay format. Lines indicate mean slopes, dotted lines give standard error. Linear regression of three biological triplicates. **C** Time-resolved emergence of plaques in HAdV-C2-dE3B-GFP-infected A549 cells treated with 1.25 μM Nelfinavir. Plaques in infected, non-treated wells shown in green, Nelfinavir-treated wells in orange and nuclei in blue. Data points represent one of eight technical replicas. Coloured vertical lines indicate means and error bars the standard deviations. **D** The inhibitory effect of Nelfinavir on HAdV-C2-dE3B-GFP spread is dependent on the amount input virus during initial infection. Number of infected, GFP-positive cells shown in green at 3 μM Nelfinavir relative to the mean infection of solvent-treated cells infected with the corresponding dosage. Total number of nuclei shown in blue, number of PI-positive dead cells in red. Note that the number of infected cells at 43 hpi is not affected by the Nelfinavir treatment. Data points represent means of four technical replicates. Dotted lines indicate standard deviation. **E** Treatment of HAdV-C2-dE3B-GFP-infected A549 cells with 1.25 μM Nelfinavir suppresses comet-shaped plaques and reveals slowly growing quasi-round plaques. Viral GFP expression levels shown as 16-color LUT. Scale bar is 1 mm. **F** Nelfinavir (1.25 μM) inhibits HAdV-C2-dE3B-GFP infection of A549 cells by slowing plaque formation. Number of infected cells and plaques per well of DMSO-treated, infected wells are shown in green, those of Nelfinavir-treated, infected wells in orange. Data points represent means of 24 technical replicates, including the example well of the micrographs shown in **D**. Error bars indicate standard deviation. Statistical significance compared to non-treated control by Kolmogorov-Smirnov test, p value < 0.0001 (****). **G** The delayed HAdV-C2-dE3B-GFP plaques in presence of 1.25 μM Nelfinavir are significantly rounder than control plaques, as indicated by Kolmogorov-Smirnov test. Data points indicate plaque regions in the well centre harbouring a single peak region. Summary of 24 technical replicates including the example well of the micrographs shown in **D**. Regions consisting of at least 5 infected cells (≥1,500 μm^2^) were considered as a plaque. Plaque morphologies in control wells could not be quantified later than 3 dpi due to rapid virus dissemination. Plaques from DMSO-treated cells 3 dpi compared to Nelfinavir-treated ones 5 dpi: approximate p value < 0.0001 (****). DMSO-treated plaques 3 dpi vs. Nelfinavir-treated plaques 6 dpi: approximate p value < 0.0001 (****). Statistical significance by Kolmogorov-Smirnov test.

We next assessed the duration of the Nelfinavir block against HAdV-C2 infection transmission. We detected strongly reduced numbers of infected nuclei and plaques in cells treated with 3 μM Nelfinavir at infection concentrations up to 100 pfu/well at 95 hpi, (Figure 3D). Remarkably, HAdV-C2-dE3B-GFP formed delayed plaques in presence of Nelfinavir, starting at 4 dpi (Figures 3E, 3F). These late plaques showed a strikingly round morphology, which was calculated to be significantly different from the comet-shaped plaques early in infection of control cells (Figure 3G). The direction of the comet tail of lytic plaques can be aligned by tilting of the incubation plate (Yakimovich et al., 2012). Thereby, the cell monolayer is positioned non-orthogonally to the vector of thermal convection flux of the liquid cell culture medium. While the direction of the comet-shaped plaques could be aligned using this method in the non-treated infections, the late Nelfinavir plaques remained mostly round (Supplementary Figures 2A-C). Moreover, there was no correlation between the size of the plaques and their roundness irrespective of Nelfinavir up to 7 dpi, demonstrating that the round plaques did not change morphology over time (Supplementary Figure 2D). Collectively, the data indicate that virus transmission in presence of Nelfinavir is not driven by the bulk current of cell free medium.

### HAdV inhibition by Nelfinavir depends on ADP

ADP is expressed at high levels late in infection and enhances cell lysis (Tollefson et al., 1992, 1996b). To test if ADP was required for Nelfinavir inhibition of lytic spread, we generated an ADP-depleted HAdV-C2-dE3B-GFP mutant, HAdV-C2-dE3B-GFP-dADP. The mutant completely lacks ADP expression, as indicated by immunofluorescence and Western blot experiments (Supplementary Figure 3A, 3B). HAdV-C2-dE3B-GFP-dADP formed particles indistinguishable from HAdV-C2-dE3B-GFP, as indicated by negative stain EM (Supplementary Figure 3C). HAdV-C2-dE3B-GFP-dADP showed a delayed onset of plaque formation by about 1 day, compared to HAdV-C2-dE3B-GFP (Figure 4A). These data are in agreement with previous kinetic studies with the ADP deletion mutant HAdV-C dl712 (Tollefson et al., 2003) (see also Supplementary Figure 3A). HAdV-C2-dE3B-GFP-dADP plaques were comet-shaped, albeit their comet-heads appeared bigger and more dense (Figure 4A). While the parental virus was highly sensitive to Nelfinavir, HAdV-C2-dE3B-GFP-dADP required much higher concentrations of the compound to show inhibition of plaque formation (Figure 4B, Supplementary Table 2). In accordance, the ADP-deleted virus induced cell death independent of Nelfinavir, unlike the ADP-expressing virus, as concluded from cell impedance measurements with xCELLigence (Figure 4C, Supplementary Figures 3D, 3E). Finally, HAdV-C2-dE3B-GFP-dADP exhibited a strongly diminished separation of anti-viral efficacy from toxicity, as indicated by reduced TI_50_ values compared to the parental virus, for example 2.1 versus 66.8 with A549 cells, 8.9 versus 61.0 with HeLa cells, and 4.6 versus 55.2 with HBEC cells (Figure 4D). These effects were in agreement with similar experiments performed with the previously described ADP-knock out mutant dl712 and the parental rec700, an HAdV-C5/2 hybrid virus (Deutscher et al., 1985; Tollefson et al., 1996b). The data are shown in (Supplementary Figures 3F to 3H). Together, the results show that the selective antiviral effects of Nelfinavir are more cell-type dependent in case of HAdV lacking ADP than in ADP-expressing viruses, and the effects are comparatively small for viruses lacking ADP.

**Figure 4.**
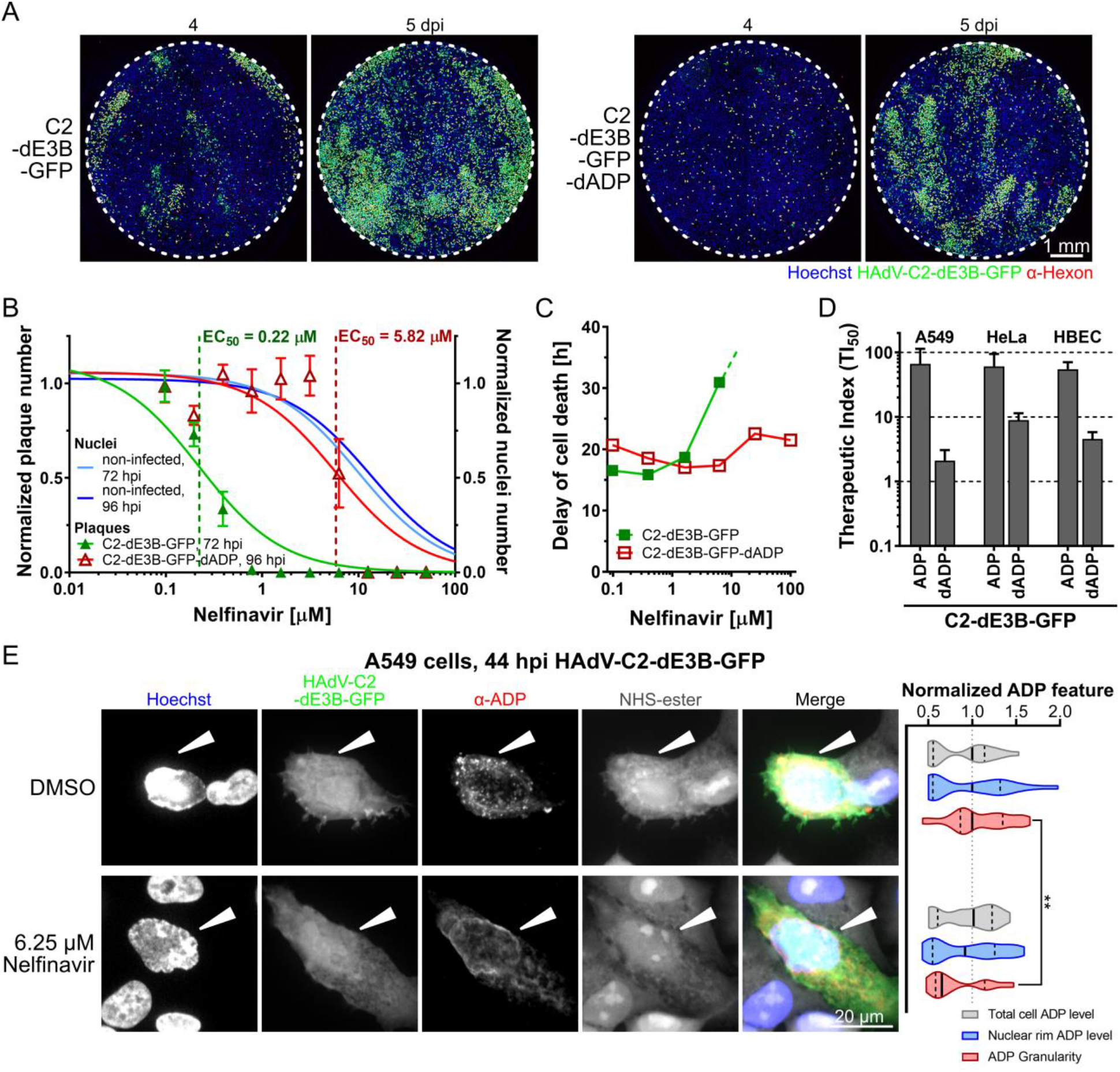
ADP contributes to the inhibitory effect of Nelfinavir against HAdV-C. **A** The deletion of ADP from HAdV-C2-dE3B-GFP delayed plaque formation in A549 cells by one day, but does not change plaque shape. Cells were infected with 1.1*10^5^ VP/ well. GFP is in green, hexon staining red, Hoechst signal of nuclei blue. Scale bar is 1 mm. **B** The deletion of ADP from HAdV-C2-dE3B-GFP reduced the anti-viral effects of Nelfinavir in A549, with EC_50_ = 5.82 compared to 0.22 μM for the parental virus. HAdV-C2-dE3B-GFP infection was quantified at 72 hpi, and 96 hpi for the ADP deletion mutant. Plaque numbers per well were normalized to the mean DMSO control and depicted as full green triangles for HAdV-C2-dE3B-GFP and empty red triangles for the ADP deletion mutant. Nuclei numbers of non-infected, treated wells were normalized to the mean DMSO control and depicted as full blue circles (72 h incubation), and empty blue circles (96 h). Data points represent means of four technical replicates. Error bars indicate standard deviation. EC_50_s were derived from non-linear curve fitting. For detailed information and statistics, see Supplementary Table 2. **C** The delay of dell death was calculated from the highest mean cell index (CI) and its half maximum for each treatment (mean of two technical replicates). HAdV-C2-dE3B-GFP data in green and dADP in red. For HAdV-C2-dE3B-GFP-infected A549 treated with 25 μM Nelfinavir, the measurement was aborted due to overgrowth causing cytotoxicity before the maximal cell index was reached. Treatment with 100 μM Nelfinavir was toxic. **D** Therapeutic index (TI_50_) derived from the ratio of Nelfinavir concentration causing 50% toxicity (TC_50_) and the concentration leading to 50% reduction in numbers of plaques per well (EC_50_). Results are shown for HAdV-C2-dE3B-GFP and -dADP in different cancer and primary cells. For detailed information and statistics, see Supplementary Table 2. **E** Representative high-magnification confocal images of HAdV-C2-dE3B-GFP-infected A549 cells 44 hpi showing the effect of Nelfinavir on ADP localization. Nuclei, shown in blue in the merged right-most panel, were stained with Hoechst. Viral GFP is displayed in green. ADP stained by immunofluorescence with a rabbit α-HAdV-C2-ADP87-101 antibody (red). Cells were stained using NHS-ester (grey scale). White arrow heads highlight infected cells. Images are max projections of 30 z planes with 0.5 μm z step, scale bar indicates 10 μm. Right: Quantification of total ADP expression (grey), ADP localization to the nuclear rim (blue) and ADP granularity (red) relative to the mean values from DMSO control cells. Data set is comprised of 20 Nelfinavir-treated infected cells, and 23 control cells. Solid line indicates median, dotted lines reflect 5-95% quantile. Significance was tested using the Kolmogorov-Smirnov test: ADP granularity p value = 0.0019 (**).

Finally, we performed immunofluorescence experiments with HAdV-C2-dE3B-GFP-infected A549 cells at 44 hpi (Figure 4E). Under non-perturbed conditions, ADP accumulated in cytoplasmic foci and the nuclear envelope. Nelfinavir treatment did not affect the overall ADP expression levels nor the amount of ADP in the nuclear periphery, including the nuclear envelope, but completely abolished the cytoplasmic ADP foci as indicated by granularity quantifications (Figure 4E, right graph). Intriguingly, Tollefson and co-workers observed earlier that ADP lacking lumenal O-glycosylation sites did not localize to large cytoplasmic granules and the corresponding HAdV-C mutant pm734.4 was non-lytic (Tollefson et al., 2003). We speculate that the localization of ADP in cytoplasmic organelles, such as Golgi compartments, where O-glycosylation occurs (Reily et al., 2019), could enhance the cell lytic function of ADP. Together, the data show that ADP is a major susceptibility factor for inhibition of HAdV-C infection spread by Nelfinavir.

### A round non-lytic plaque phenotype in HAdV-C infection

Viruses are transmitted between cells by three major mechanisms, cell-free through the extra-cellular medium, directly from cell-to-cell, or in an organism by means of infected motile cells or fluid flow in blood or lymphoid vessels. This can result in far-reaching or mostly local virus dissemination (for a simplified cartoon, see Figure 5A). In cell culture, HAdV-C transmission from a lytic infected cell (staining PI-positive) yields comet-shaped infection foci due to convective passive mass flow in the cell culture medium (Yakimovich et al., 2012, 2015), consistent with lytic HAdV-C infection (Doronin et al., 2003; Tollefson et al., 1996b). In accordance, neutralizing antibodies against HAdV-C2 added to the cell culture medium suppressed the comet-shaped plaques of HAdV-C2-dE3B-GFP, and yielded confined, predominantly round-shaped infection foci 4 dpi, akin to Nelfinavir-treated infections (Figure 5B).

**Figure 5.**
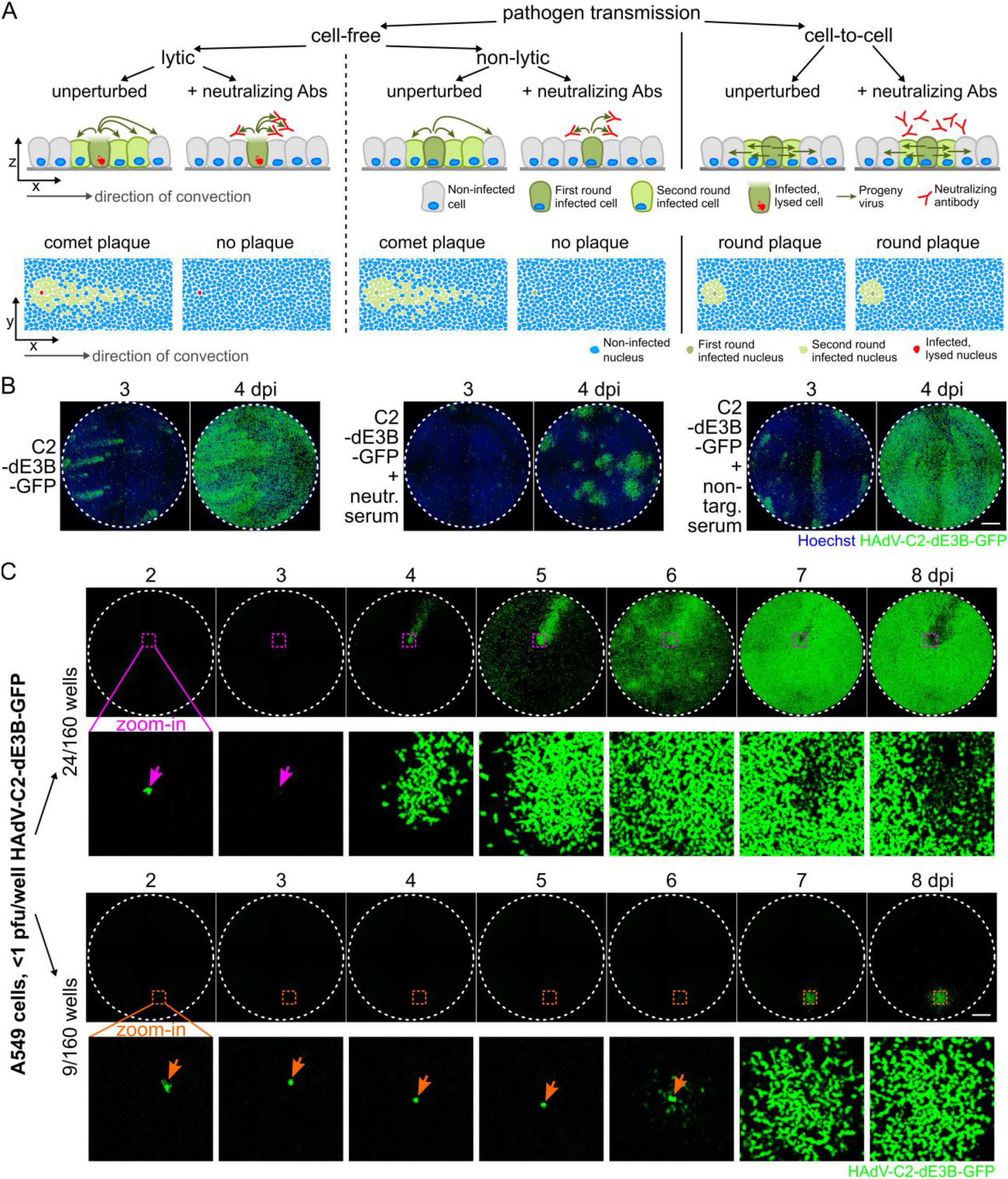
Round plaque phenotypes in presence of neutralising anti-HAdV-C2 antibodies and in unperturbed HAdV-C2 infections. **A** Schematic overview of pathogen transmission routes in cell cultures. Cell lysis kills the donor cell and releases progeny, while non-lytic egress maintains the infected donor cell. In both cases, convection in the media leads to long-distance, comet-shaped plaques. Cell-free virus transmission is susceptible to neutralizing antibodies. In contrast, direct cell-to-cell pathogen spread from persisting first round infected host cells causes symmetric, small-growing, dense round plaques, which cannot be inhibited by neutralizing antibodies presenting the extracellular medium. Non-infected cells are shown in grey with blue nuclei. First round infected cells are shown in dark green, red nuclei contain a ruptured nuclear envelope. Second round infected cells are shown in light green. Grey arrow represents direction of convective flow. Axes indicate side or top-down view. **B** Inhibition of cell-free HAdV-C2-dE3B-GFP transmission by an anti-HAdV-C2/5-neutralizing serum added to the medium at 1:10. Nuclei are shown in blue, viral GFP in green. **C** Infection of A549 cells with limiting amounts of HAdV-C2-dE3B-GFP (<1 pfu/ well, 9-75 VP/ well) in 160 wells gives rise to 33 single plaques / well. Twenty four wells contained GFP-positive comet-shaped plaques (upper panel), and nine developed delayed round plaques (lower panel). Dashed coloured squares indicate magnified regions of first-round infected cell below. Infected cell leading to comet-shaped plaque (upper panel, pink arrow) lyses at 3 dpi as indicated by loss of GFP signal. Infected cell giving rise to round plaque (lower panel, orange arrow) remains GFP-positive. Scale bar is 1 mm.

To test if round-shaped infection foci (plaques) occurred in regular HAdV-C2-dE3B-GFP infections, we analysed A549 cells infected with less than 1 plaque forming unit(s) (pfu) per well in 160 wells of 96-well formats up to 8 dpi. Thirty three wells developed a single plaque, and 24 of them contained fast emerging comet-shaped plaques (Figure 5C, upper panel), while nine developed delayed round plaques starting 6 dpi (Figure 5D, lower panel). The originally infected cell (indicated by the pink arrow), which gave rise to the comet-shaped plaque, disappeared between 2 and 3 dpi. In contrast, the infected cell giving rise to the round plaque (orange arrow) remained GFP-positive and apparently viable when the surrounding cells were infected. These data suggest that HAdV-C2 utilizes both lytic and non-lytic transmission, the former involving cell-free transmission, and the latter cell-associated transmission.

### Nelfinavir has a broad anti-HAdV spectrum

We finally assessed the inhibition breadth of Nelfinavir against various HAdV types from species A, B, C and D in different human cell lines, as well as mouse adenovirus (MAdV) 1 and 3 in mouse rectum carcinoma CMT93 cells. To balance statistical significance and automated plaque segmentation, we first determined the optimal amount of inoculum and duration of infection for each virus and cell line. The resulting TI_50_ values of Nelfinavir were heterogeneous for different HAdV types, as determined in A549 cells (Figure 6A, for details see Supplementary Table 2). While all the tested HAdV-C types as well as HAdV-B14 showed high TI_50_s (>10) ranging from 12.22 (HAdV-C1) to 71.09 (HAdV-C2). Members of HAdV species A, D and most of the HAdV-B types showed intermediate (2 - 10) to low Nelfinavir susceptibility (<2), notably HAdV-B7 and B11 with TI_50_<1. MAdV-1 and 3 also showed low susceptibility. Noticeably, a high susceptibility of HAdV-C was consistently observed in human lung epithelial carcinoma (A549) cells, human epithelial cervix carcinoma (HeLa) cells, immortalized primary normal human corneal epithelial (HCE) cells as well as normal human bronchial epithelial (HBEC) cells. The corresponding TI_50_ values were in the same range as for herpes simplex virus (HSV) 1, for which Nelfinavir was reported to be an egress inhibitor (Gantt et al., 2011, 2015; Kalu et al., 2014).

**Figure 6.**
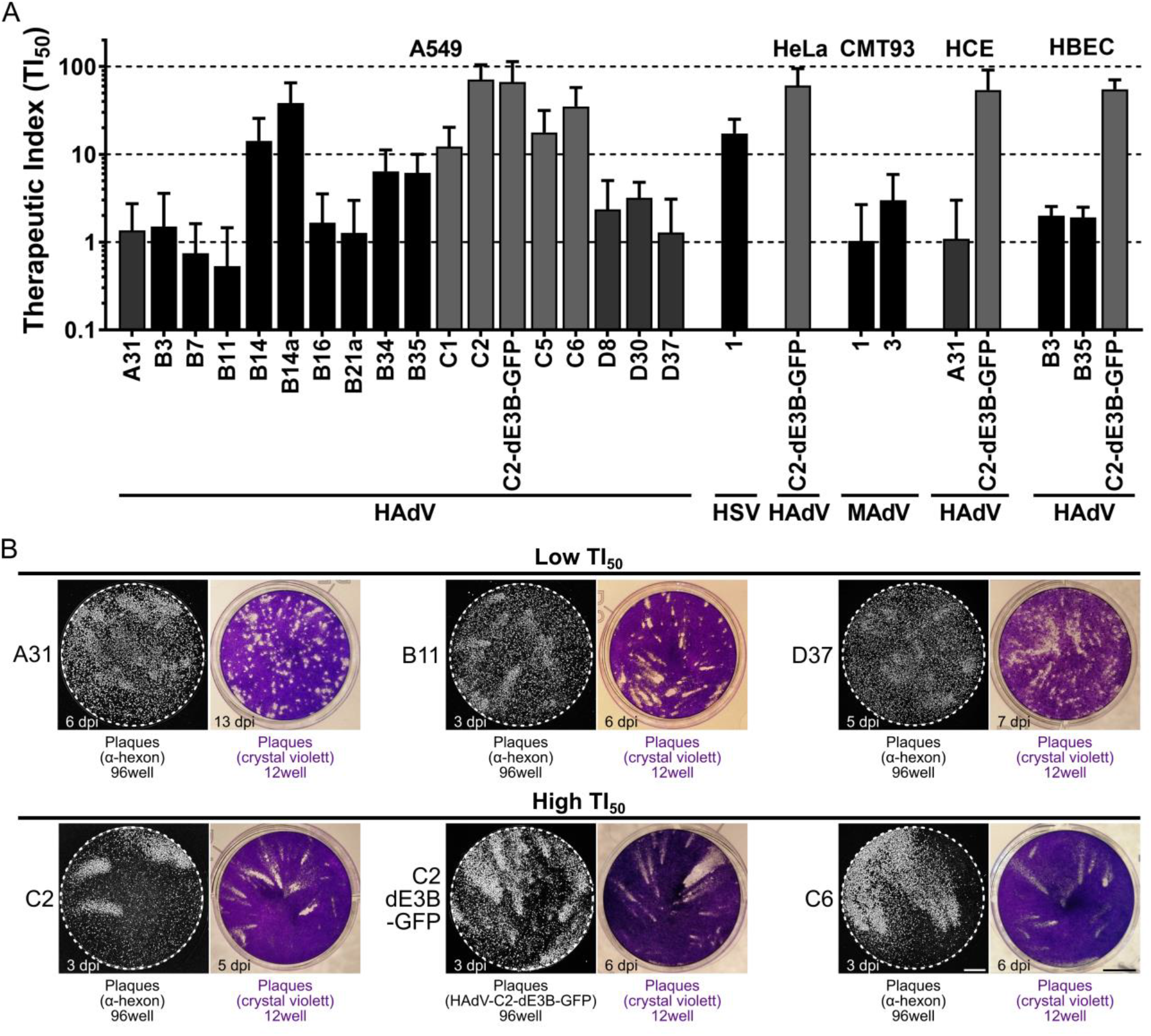
Susceptibility of HAdV to Nelfinavir correlates with plaque shape. **A** Therapeutic index (TI_50_) calculated as Nelfinavir concentration causing 50% toxicity (TC_50_) divided by the concentration leading to 50% reduction in numbers of plaques per well (EC_50_) for different HAdV, mouse adenoviruses (MAdV) and herpes simplex virus 1 (HSV-1) in different cancer and primary cell lines. For detailed information and statistics, see Supplementary Table 2. **B** Representative microscopic and macroscopic plaque morphologies of Nelfinavir-sensitive and insensitive HAdV types. Grey scale images show plaques based on epifluorescence microscopy of hexon immunostaining or GFP expression in a 96-well of A549 cells infected with the indicated HAdV type. Scale bar is 1 mm. Coloured images show infection-induced cytotoxicity yielding plaques, as visualized by crystal violet staining in wells of a 12-well dish of A549 cells infected with the indicated HAdV type. Scale bar is 5 mm.

We finally examined the plaque morphologies in non-perturbed infections by immunofluorescence staining of the late proteins VI and hexon, as well as macroscopic analyses of crystal violet stained dishes for classical plaques (Figure 6B). Viruses that were highly susceptible to Nelfinavir (exhibiting high TI_50_ values) formed exclusively comet-shaped plaques. Viruses with low TI_50_ values, such as A31, B11 or D37 had a high fraction of round plaques, even when infected with more than 1 pfu / well. This demonstrates that the slowly growing round infection foci observed in fluorescent microscopy gave similarly shaped lesions due to cytotoxicity, akin to the lytic comet-shaped foci. We conclude that HAdV types employ lytic cell-free and non-lytic cell-to-cell transmission modes and give rise to different plaque phenotypes.

## Discussion

A phenotypic screen of the PCL identified Nelfinavir as a potent, post-exposure inhibitor of HAdV-C2-dE3B-GFP plaque formation in cell culture (Georgi et al., 2020). Nelfinavir is a non-nucleoside class inhibitor against a range of HAdV types. Surprisingly, we found Nelfinavir to inhibit HAdV infection, although Nelfinavir was previously classified as inactive against HAdV-C based on replication assays (Gantt et al., 2011). It is the off-patent FDA-approved active pharmaceutical ingredient of Viracept. Nelfinavir was originally developed as an inhibitor against the HIV aspartyl protease. It is orally bioavailable, with an inhibitory concentration in the low nanomolar range (Kaldor et al., 1997; Moyle et al., 1998). Nelfinavir inhibits the replication of enveloped viruses, including SARS coronavirus (Yamamoto et al., 2004), hepatitis C virus (Toma et al., 2009) as well as α-, β- and γ-herpes viruses (Gantt et al., 2011). In the case of the α-herpes virus HSV-1, Nelfinavir inhibits the envelopment of the capsid with cytoplasmic membranes (Gantt et al., 2011, 2015; Kalu et al., 2014). Nelfinavir was reported to inhibit the activity of regulatory proteases in the Golgi, the growth of cancer cells and to induce a wealth of other effects, including autophagy, ER stress, the unfolded protein response, and apoptosis (Caron et al., 2003; Chow et al., 2006; Gills et al., 2007; Guan et al., 2011, 2012, 2015; Yang et al., 2005) (reviewed in Bernstein and Dennis, 2008; Brüning et al., 2010; Chow et al., 2009; Koltai, 2015; Shim and Liu, 2014).

Here, we demonstrate that Nelfinavir inhibits the egress of HAdV particles without perturbing other viral replication steps including entry, assembly and maturation. Morphometric analyses of the fluorescent plaques indicated that HAdV-C propagates by two distinct mechanisms, lytic and non-lytic. Lytic transmission led to comet-shaped convection driven plaques, whereas non-lytic transmission gave rise to symmetric round-shaped plaques. Nelfinavir specifically suppressed the lytic spread of HAdV, most prominently the HAdV-C types and B14, but not other HAdV, such as A31 or D37. Incidentally, HAdV-C and B14 replicate to considerable levels in Syrian hamsters, whereas other HAdV types do not (Radke et al., 2016; Tollefson et al., 2017; Wold et al., 2019). We infer that lytic infection could be a pathogenicity driver, at least in the hamster model.

The molecular mechanisms underlying cell lysis in AdV infection are not well understood, largely due to the lack of specific assays and inhibitors. Single cell analyses combined with machine learning start to identify specific features of lytic cells, such as increased intra-nuclear pressure compared to non-lytic cells (Andriasyan et al., 2019). The lysis induced by HAdV was suggested to involve caspase-dependent functions, and necrosis-like features (Abou El Hassan et al., 2004; Yun et al., 2005; Zou et al., 2004). The best characterized factor in HAdV cell lysis is ADP, a small membrane protein encoded in HAdV-C (Davison et al., 2003a, 2003b; Robinson et al., 2013). ADP-deletion mutants show delayed onset of plaque formation (Tollefson et al., 1996a, 1996b). Lysis is enhanced by increased ADP levels and tuned by post-translational ADP processing (Doronin et al., 2003; Tollefson et al., 1996a, 1996b). ADP has a single signal/anchor sequence, and its lumenal domain is N- and O-glycosylated. The N-terminal segment is cleaved off in the Golgi lumen, and the membrane-anchored ADP localizes to the inner nuclear membrane (Scaria et al., 1992; Tollefson et al., 1996b, 2003). Interestingly, two cysteine residues in the cytoplasmic domain adjacent to the transmembrane segment are palmitoylated (Tollefson et al., 2003) (Hausmann et al., 1998). S-palmitoylation is known to support anchorage and sorting of host and viral membrane proteins. Accordingly, S-palmitoylation in the Golgi facilitates protein oligomerization, virion assembly and entry, as shown for structural proteins of enveloped viruses, including SARS-CoV-1 S, vesicular stomatitis virus G, sindbis virus E2, influenza virus HA, respiratory syncytial virus F, or rubella virus E1 and E2, as well as viroporin-mediated membrane permeabilization, including mouse hepatitis virus E protein, SARS-CoV-1 E protein and sindbis virus 6K. For reviews, see (Blaskovic et al., 2013; Veit, 2012).

Conspicuously, the cell lysis defective HAdV mutant pm734.4 encodes a C2 mutant ADP with two point mutations in the transmembrane domain, C_53_R and M_56_L (Tollefson et al., 2003). The mutant ADP localizes to the ER and the nuclear envelope, but not the Golgi, unlike the parental wild type virus rec700. The localization of the pm734.4 ADP is akin to the localization of HAdV-C2 ADP in Nelfinavir-treated cells, which resist lysis and lack ADP localization in the Golgi. We speculate that the palmitoylation of ADP in the Golgi is crucial for ADP to enhance the rupture of the nuclear membrane in lytic HAdV-C egress. Nelfinavir may interfere with ADP palmitoylation either by inhibiting a palmitoyl-acyltransferase or by dispersing the donor substrate for protein palmitoylation, palmitoyl-coenzyme A (Blaskovic et al., 2013). Remarkably, Nelfinavir has a high logP value, 4.1 to 4.68 (Longer et al., 1995; Tetko et al., 2005), and partitions into lipophilic domains of the cell, including membranes. This is akin to another lipophilic drug with pleiotropic effects, the anti-viral and anti-helminthic compound Niclosamide, which is a weak acid and acts as a protonophore extracting protons from acidic organelles, and thereby inhibits virus entry and uncouples mitochondrial proton gradients (Fonseca et al., 2012; Jurgeit et al., 2012).

We noticed that ADP is not the sole lysis factor of HAdV. HAdV types lacking ADP, such as B types, also release their progeny by lysis, albeit with efficacies that vary depending on the cell type (Baker et al., 2019; Chen et al., 2011a; Uchino et al., 2014). This is in agreement with the observation that HAdV types of the A, B and D species form comet-shaped plaques, and that ADP-deleted HAdV-C2 lyse the host cell, and form comet-shaped plaques, albeit delayed and with lower efficacy than ADP-containing rec700 or HAdV-C2-dE3B-GFP.

In addition to providing a new inhibitor of lytic HAdV propagation, Nelfinavir revealed an alternative non-lytic HAdV transmission pathway, which gives rise to slow-growing symmetrical plaques. This non-lytic pathway exists in unperturbed cells, but is camouflaged by the rapid and far-reaching lytic infection. Non-lytic egress from the nucleus bypasses the nuclear envelope and the plasma membrane. We speculate that the non-lytic pathway involves sorting of HAdV particles to membrane sites where outward budding and scission occur. HAdV budding through the nuclear envelope could involve the WASH complex, akin to nuclear release of large RNPs in Drosophila, and perhaps similar to HSV budding (Hagen et al., 2015; Verboon et al., 2019). Cytoplasmic membrane budding could be enhanced by the ESCRT complex, which is known to release enveloped viruses, such as HIV, and also facultative-enveloped viruses, such as hepatitis A virus (Feng et al., 2013; Hurley, 2015; Lippincott-Schwartz et al., 2017). Alternatively, autophagy could sequester virions from the nucleus and upon fusion with the plasma membrane release virions from infected cell.

In conclusion, our work opens new therapeutic options for treating adenovirus disease, including acute and persistent infections. For example, HAdV-C persists in lymphocytes, which resist lytic infection, but also in epithelial cell lines under the repression of interferon and activation of the unfolded protein response sensor IRE-1a (Garnett et al., 2002; Kosulin et al., 2016; Murali et al., 2014; Prasad et al., 2014, 2020b; Zheng et al., 2016). Nelfinavir might be considered for anti-HAdV therapy, for example prophylactically in hematopoietic stem cell recipients, whose life is threatened by reactivation of HAdV-C (Hiwarkar et al., 2018; Lion, 2019; Lynch and Kajon, 2016).

## Contributions

UFG, VA and AY conceived the project. FG, VA, RW, LY, MG and NM performed experiments. FG, VA, RW, LM, FK, VP, AY, GT, UFG analysed data. SH contributed essential viruses. FG and UFG wrote the manuscript.

## Acknowledgements

We thank Anja Ehrhardt for providing the HAdV-C-neutralizing dog serum. William SM Wold and Anne Tollefson kindly provided the α-ADP antibodies and ADP mutant viruses. Albert Heim provided HAdV-B14 and 21, Maarit Suomalainen provided HAdV-C2 and C5, discussions and advice. Yohei Yamauchi contributed Influenza A virus H1N1 virus and Cornel Fraefel the HSV-1-CMV-GFP. Vibhu Prasad kindly supported the impedance experiments. We are grateful for scientific discussions with Ivo Sbalzarini. We thank the Center for Microscopy and Image Analysis (ZMB) at UZH providing the EM instrumentation. We thank the members of the Greber lab for constructive discussions.

## Abbreviations

ADP: adenovirus death protein
AdV: adenovirus
CAR: coxsackievirus adenovirus receptor
CMV: cytomegalovirus
DFT: 3′-Deoxy-3′-fluorothymidine
DMEM: Dulbecco’s Modified Eagle medium
dpi: day(s) post infection
EC_50_: 50% effective concentration
GFP: green fluorescent protein
HAdV: human adenovirus
HIV: human immunodeficiency virus
hpi: hour(s) post infection
HSV: herpes simplex virus
kbp: kilo base pairs
MAdV: mouse adenovirus
o/n: over night
ORF: open reading frame Nelfinavir Mesylate Nelfinavir
pBI: pBluescript
PCL: Prestwick Chemical Library
PFA: para-formaldehyde
pfu: plaque forming unit(s)
pi: post infection
PI: propidium iodide
RT: room temperature
SE: standard error
SD: standard deviation
TC_50_: 50% toxic concentration
TI: therapeutic index
ts: temperature-sensitive
VP: viral particles
wt: wild type

## Supplementary Figures

**Supplementary Figure 1.**
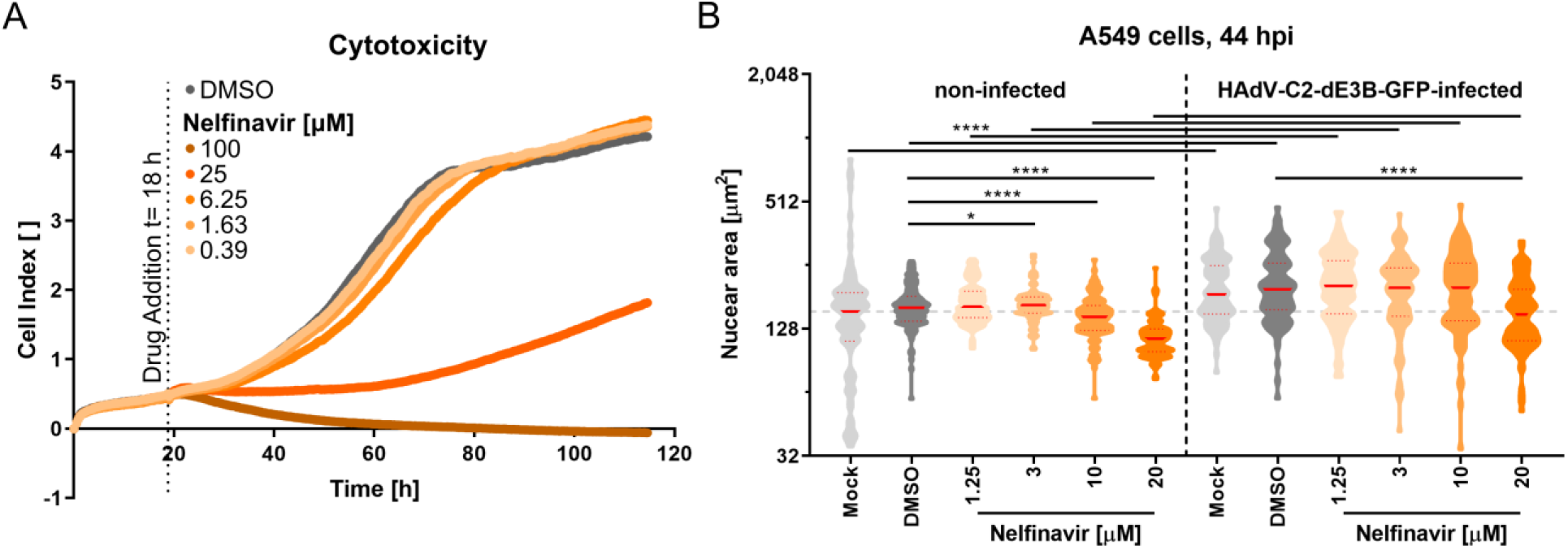
Nelfinavir exhibits low toxicity. **A** Cell index (CI) profiles of uninfected A549 shows little signs of toxicity up to 6.25 μM of Nelfinavir during more than 4 days. Impedance was recorded every 15 min using xCELLigence. Vertical dotted line shows the time of drug addition. **B** Nelfinavir does not affect infection-induced nuclear swelling in HAdV-C2-dE3B-GFP-infected A549 cells at 44 hpi. Each violin symbol represents the areas of 99 nuclei from four technical replicates. Significance was tested using the Kolmogorov-Smirnov test, p value ≤0.05 (*), ≤0.0001 (****). Difference is not significant, where none is indicated. Solid red lines indicate median, dotted red lines mark 5-95% quartile. Horizontal dotted grey line at median of untreated, uninfected nuclei. Note that Nelfinavir induced the shrinkage of the nuclei at 20 μM in both infected and uninfected cells, indicative of toxicity at high concentrations.

**Supplementary Figure 2:**
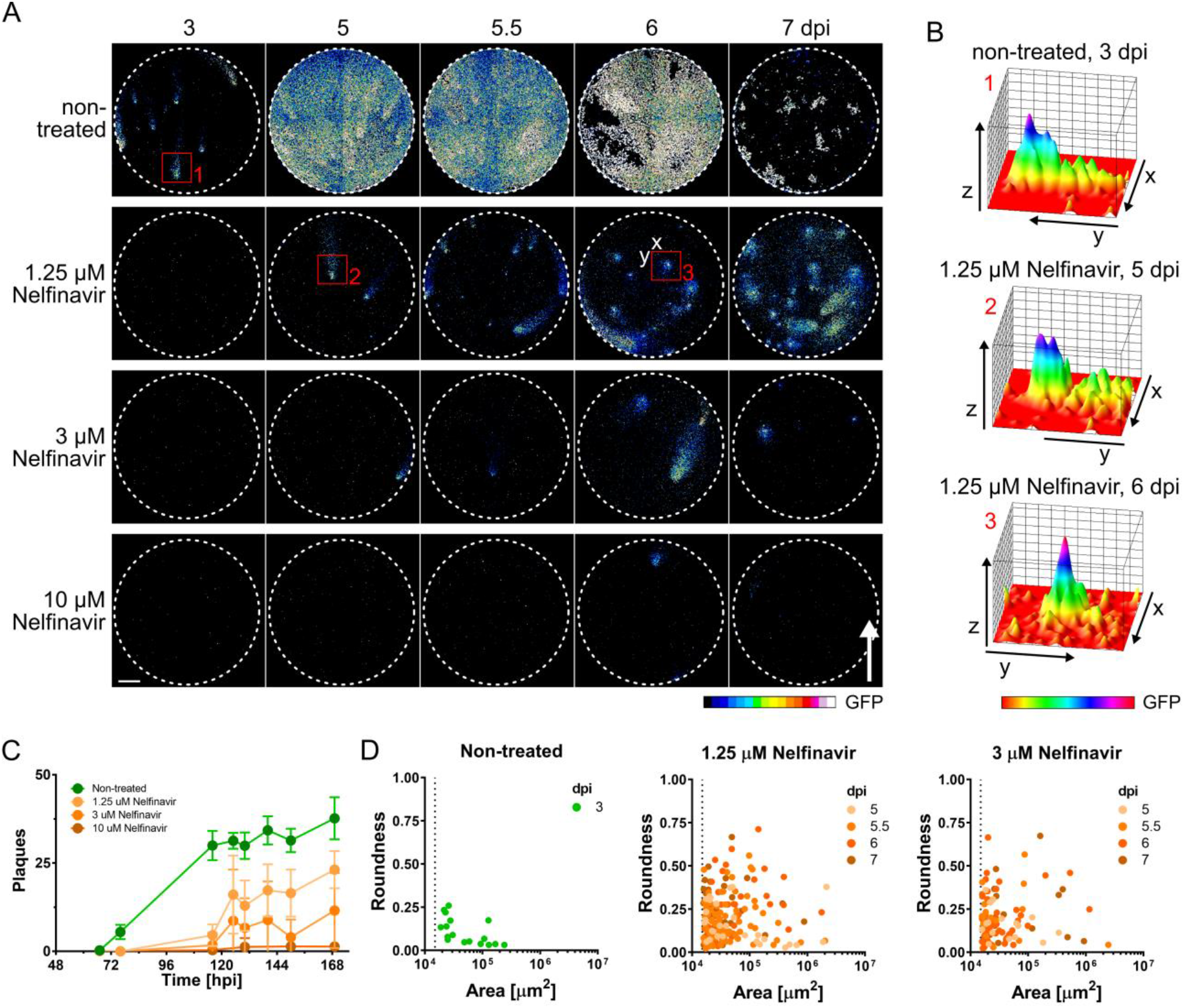
Characterization of round plaque phenotypes revealed by Nelfinavir. **A** Plate tilting to direct convection of medium fluids does not affect the spatio-temporal spread pattern of HAdV-C2-dE3B-GFP in presence Nelfinavir. Nelfinavir delays the formation and number of plaques in a concentration-dependent manner. Early plaques in absence of Nelfinavir show comet-shaped morphologies (red squares 1 and 2), while late plaques in presence of Nelfinavir appear round as exemplified by red square 3. GFP intensity is shown as 16-color LUT. White arrow indicates uphill flow of convection, and direction of elongation of comet plaques. Scale bar is 1 mm. **B** Three-dimensional topological views of plaque morphologies generated by depicting the viral GFP expression level along the z-axis. Red numbers correspond to regions of interest (ROIs) 1 to 3, indicated by red squares in **A**. x- and y-axes are oriented as labelled in **A**. GFP intensity is shown as rainbow colour LUT, indicating the original of plaque formation where GFP intensity peaks (purple). **C** Delay in plaque formation by Nelfinavir over time. Data points represent means of 12 replicates from two experiments, including the example well micrographs shown in **A**. Error bars indicate standard deviations. DMSO-treated infected wells are shown in green, infected wells treated with 1.25, 3 or 10 μM Nelfinavir are shown in shades of orange. **D** Morphological analysis of plaque roundness compared to size over the course of Nelfinavir treatment at 1.25 and 3 μM at different times of infection (shades of orange). Data points indicate centre well plaque regions harbouring a single peak region from 12 well dishes and two experiments including the example well micrographs shown in **A**. Plaque morphologies in control wells could not be quantified later than 3 dpi, due to extensive virus dissemination. Regions consisting of at least five infected cells (≥1,500 μm^2^ indicated by the dotted vertical lines) were considered as plaque.

**Supplementary Figure 3.**
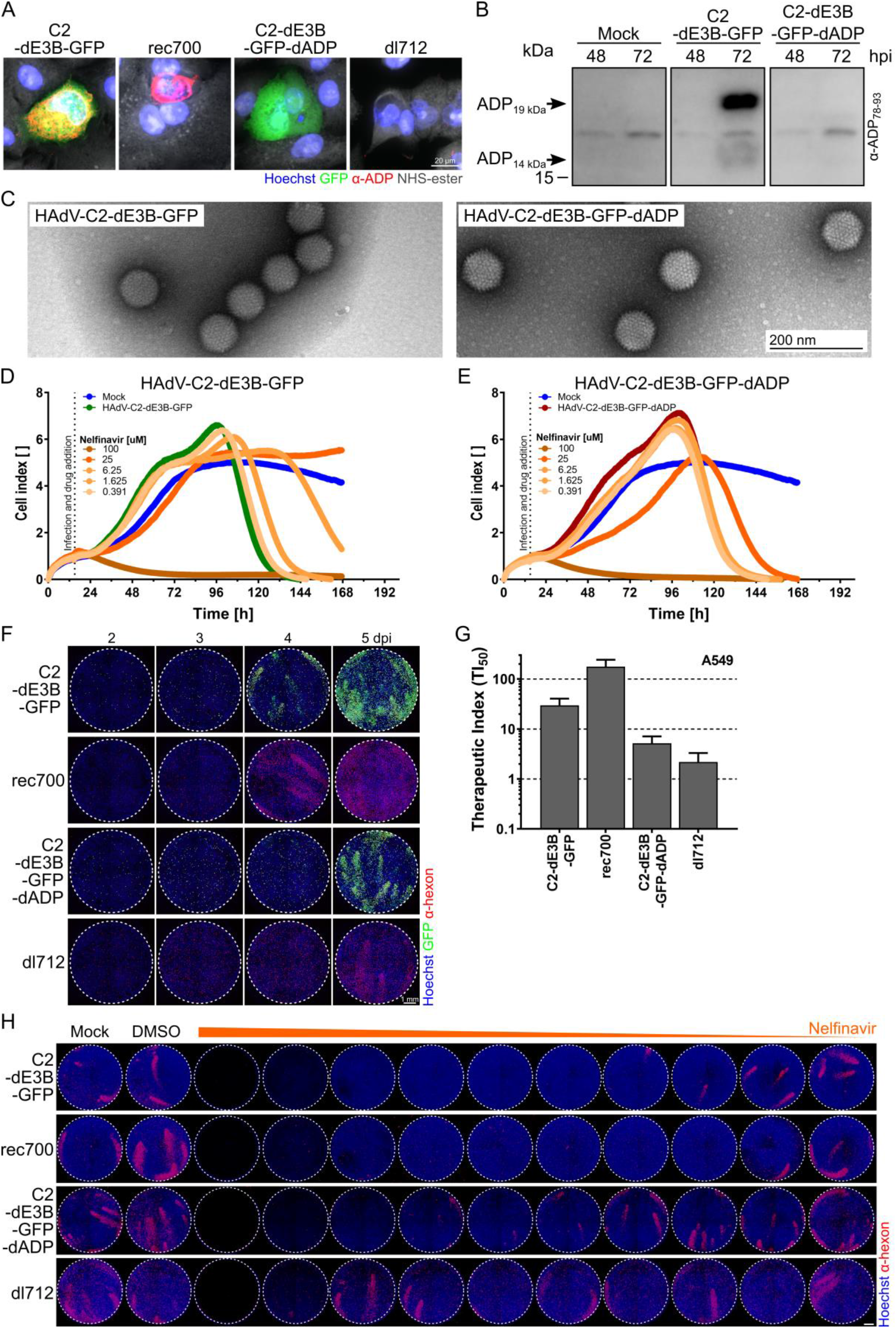
The mode of action of Nelfinavir is ADP-dependent irrespective of E3B deletion. **A** Lack of ADP in HAdV-C2-dE3B-GFP-dADP infected cells 44 hpi shown by indirect immunofluorescence. HAdV-C2-dE3B-GFP-dADP infected cells compared to dl712, the ADP-deleted HAdV-C5/2 rec700 mutant. Nuclei are shown in blue, GFP signal in green, ADP in red and cytosol staining in grey. Scale bar is 20 μm. **B** Western blot analyses of HAdV-C2-dE3B-GFP-dADP compared to HAdV-C2-dE3B-GFP infected cells 48 and 72 hpi. Full length ADP runs with an apparent mass of 19 kDa, the cleaved 14 kDa form of ADP at 16 kDa apparent mass. **C** Negative stain EM micrographs of purified HAdV-C2-dE3B-GFP and HAdV-C2-dE3B-GFP-dADP particles. Scale bar indicates 200 nm. **D, E** Mean cell index profiles from impedance measurements of infected A549 cells performed in two technical replicates indicate cytopathic effects. Nelfinavir inhibits cytotoxicity in DMSO-treated HAdV-C2-dE3B-GFP infection (**C**, green profile), but not in cells infected with the ADP-deletion mutant (**D**, red profile). Impedance was recorded every 15 min using xCELLigence. Each point represents the average value from two replicates with standard deviations. The time on the x-axis indicates hours after cell seeding. Vertical lines show the time of infection and drug addition. Profiles of non-infected cells are shown in blue. The concentrations of Nelfinavir are represented by different shades of orange. **F** Plaque formation in ADP-deleted HAdV-C2-dE3B-GFP and dl712 compared to HAdV-C2-dE3B-GFP and rec700 in A549 is delayed by 1 day, but plaque shapes are not affected at the indicated time points. Cells were infected with 1.1*10^5^ VP / well. Nuclei signal shown in blue, viral GFP in green and hexon staining in red, scale bar is 1 mm. Dotted line indicates well outline. **G** Therapeutic index (TI50), based on images shown in **H** and three additional technical replicates for HAdV-C harbouring ADP (HAdV-C2-dE3B-GFP and rec700) compared to ADP-depleted HAdV-C (HAdV-C2-dE3B-GFP-dADP and dl712). Plaque quantification was based on hexon staining. For detailed information and statistics, see Supplementary Table 2. **H** Representative well images of hexon-stained HAdV-C infection of A549 cells treated with 50 to 0.1 μM Nelfinavir (from left to right, indicated by orange triangle). Cells were fixed at 96 hpi (HAdV-C2-dE3B-GFP and rec700) or 112 hpi (HAdV-C2-dE3B-GFP-dADP and dl712). Nuclei signal is shown in blue, hexon staining in red. Scale bar is 1 mm. Dotted line indicates well outline.

## Supplementary Tables

**Supplementary Table 1.**
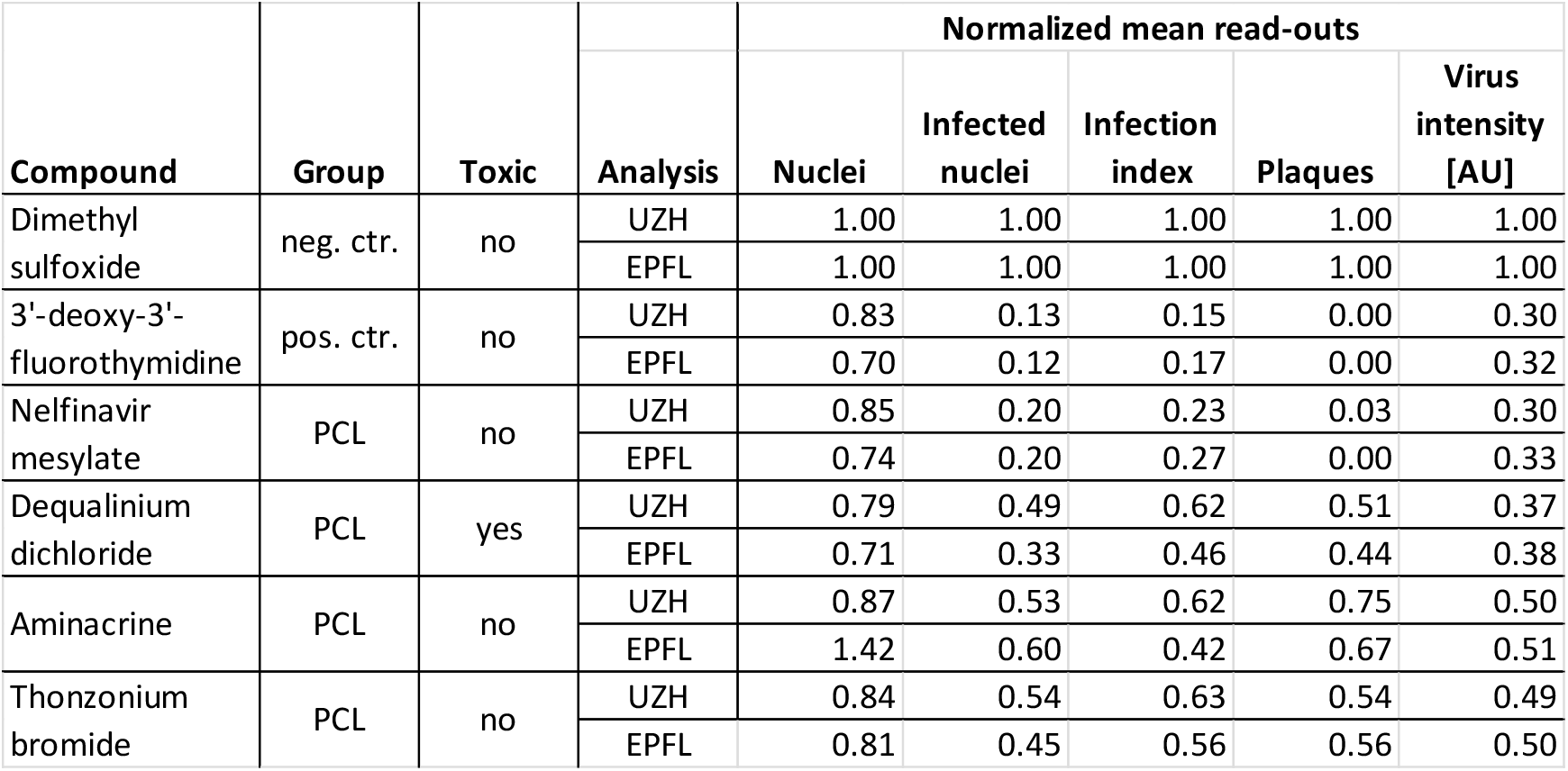
Summary of top hits identified in the HAdV AntiVir screen. An accompanying manuscript (Georgi et al., 2020) tested the Prestwick Chemical Library (PCL) for potentially repurposable inhibitors of HAdV infection. The data was acquired and analysed by two independent research teams at UZH and EPFL. The mean infection scores normalized to the negative control (neg. ctr.) obtained by both teams are listed for the four top hits. 3’-deoxy-3’-fluorothymidine (DFT) was used as positive control (pos. ctr.). Toxicity in absence of infection was tested using the same experimental outline.

| Compound | Group | Toxic | Analysis | Normalized mean read-outs |  |  |  |  |
| --- | --- | --- | --- | --- | --- | --- | --- | --- |
|  |  |  |  | Nuclei | Infected nuclei | Infection index | Plaques | Virus intensity [AU] |
| Dimethyl sulfoxide | neg. ctr. | no | UZH | 1.00 | 1.00 | 1.00 | 1.00 | 1.00 |
|  |  |  | EPFL | 1.00 | 1.00 | 1.00 | 1.00 | 1.00 |
| 3'-deoxy-3'-fluorothymidine | pos. ctr. | no | UZH | 0.83 | 0.13 | 0.15 | 0.00 | 0.30 |
|  |  |  | EPFL | 0.70 | 0.12 | 0.17 | 0.00 | 0.32 |
| Nelfinavir mesylate | PCL | no | UZH | 0.85 | 0.20 | 0.23 | 0.03 | 0.30 |
|  |  |  | EPFL | 0.74 | 0.20 | 0.27 | 0.00 | 0.33 |
| Dequalinium dichloride | PCL | yes | UZH | 0.79 | 0.49 | 0.62 | 0.51 | 0.37 |
|  |  |  | EPFL | 0.71 | 0.33 | 0.46 | 0.44 | 0.38 |
| Aminacrine | PCL | no | UZH | 0.87 | 0.53 | 0.62 | 0.75 | 0.50 |
|  |  |  | EPFL | 1.42 | 0.60 | 0.42 | 0.67 | 0.51 |
| Thonzonium bromide | PCL | no | UZH | 0.84 | 0.54 | 0.63 | 0.54 | 0.49 |
|  |  |  | EPFL | 0.81 | 0.45 | 0.56 | 0.56 | 0.50 |

**Supplementary Table 2.**
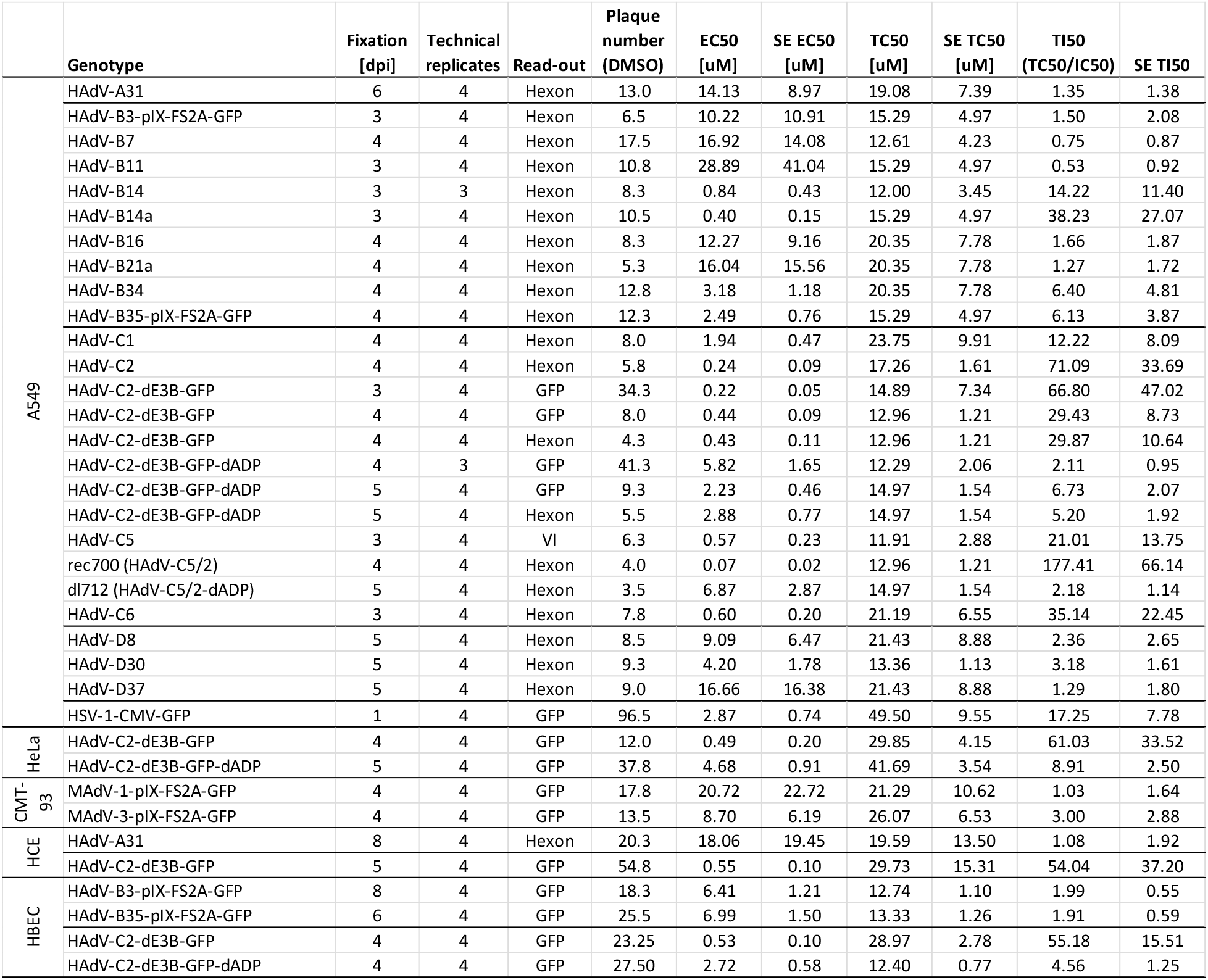
Statistical information on the TI_50_ of Nelfinavir against different viruses in different cell lines. Indicated cell lines are infected with the indicated pfu/ well virus and fixed at the indicated time post infection. Plaque numbers were quantified based on viral GFP expression or HAdV hexon immunofluorescence staining as indicated. EC_50_ indicates effective concentration 50%, Nelfinavir concentration leading to 50% reduction in plaque number/ well). TC_50_ refers to toxic concentration 50%, Nelfinavir concentration causing 50% reduction in nuclei number/ well. The therapeutic index, TI_50_, is the TC_50_/ EC_50_ ratio. All values are means of the indicated number of technical replicates. SE gives standard error. Plaque numbers were quantified using Plaque2.0. Nuclei numbers were quantified based on Hoechst signal using CellProfiler. Non-linear regression was performed in GraphPad.

## Supplementary Methods

### Generation of HAdV-C2-dE3B-GFP-dADP

As a first, a pKSB2-based bacterial artificial chromosome vector pKSB2-AdV-C2-dE3B_GFP carrying the genome of HAdV-C2-dE3B-GFP (Yakimovich et al., 2012) was generated starting with pKSB2-AdV-C2-LARAzeo containing left and right terminal HAdV-C2 fragments. To generate pKSB2-AdV-C2-LARAzeo, two PCR-generated fragments were first cloned into pBluescript (pBl). The first fragment encompassed 853 bp of the HAdV-C2 left end sequence and was PCR-amplified using the forward primer 5’-ataagaatGCGGCCGCTAGGGATAACAGGGTAATcatcatcataatataccttattttgg-3’ inserting the restriction sites NotI and I-SceI, and the reverse primer 5’-CTCTCTACTAGTAATAAGTCAATCCCTTCCTGC −3’ inserting the restriction site SpeI. HAdV-C2-dE3B-GFP genomic DNA isolated from infected A549 cells was used as template. The NotI and SpeI restriction sites were used to clone the left arm fragment into pBl. The second fragment encompassed 853 bp of the HAdV-C2 right end sequence and was PCR-amplified using the forward primer 5’-agagagACTAGTaaaaacatttaaacattagaagcctg-3’ adding the restriction sites SpeI and the reverse primer 5’-gcgcaagcttATTACCCTGTTATCCCTAcatcatcataatataccttattttgg-3’ adding the sites I-SceI and HindIII. This fragment was cloned into pBl-AdV2-C2-LA by SpeI and HindIII restriction sites. A SpeI fragment containing the zeocine resistance marker from pcDNA3.1 zeo (Invitrogen) and generated by PCR using the forward 5’-GACTAGTTTTTCG GATCTGATCAGCACG-3’ and reverse primer 5-GACTAGTGGAAAACGATT CCGAAGCCC-3’ was cloned into the SpeI of pBl-AdV-C2-LARA, connecting the two HAdV-C2 arms and resulting in pBl-AdV-C2-LARAzeo. In order to transfer the AdV-C2-LARAzeo cassette to the BACmid pKSB2, the NotI-HindIII fragment containing this sequence was ligated with the NotI-HindIII-restricted pKSB2 vector. Colonies containing pKSB2-AdV-C2-LARAzeo were selected using chloramphenicol and zeocin at concentrations of 10 μg / ml and 25 μg / ml, respectively. In order to generate pKSB2-AdV-C2-dE3B_GFP, homologous recombination was performed in SW102 bacteria using AatII and ApaLI-restricted pKSB2-AdV-C2-LARAzeo and HAdV-C2-dE3B-GFP genomic DNA isolated from infected A549 cells.

As a second, HAdV-C2-dE3B-GFP-dADP, was generated using two recombineering steps. In a first, the galK cassette was introduced into pKSB2-AdV-C2_dE3B_GFP to replace the ADP sequence. The GalK cassette was amplified using the forward primer 5’-ACTGCAAATTTGATCAAACC CAGCTTCAGCTTGCCTGCTCCAGAG cctgttgacaattaatcatcggca-3’ and the reverse primer 5’-GAACTAATGACCCCGTAATTGATTACTATTAATAA CTAGTCTCATctcagcactgtcctgctcctt −3’ introducing 45 nucleotides of flanking sequences. Subsequently, the GalK sequence was replaced with a dsDNA of the sequence actgcaaatttgatcaaacccagcttcagcttgcctgctccagagatgagactagttattaatagtaatcaattacggggtcattagttc resulting in deletion of ADP.

To generate infectious virus, circular pKSB2-AdV-C2_dE3B_GFP_dADP was transfected in human 911 cells stably expressing I-SceI endonuclease (Ibanes and Kremer, 2013) using the jetPEI transfection reagent (Polyplus transfection, Illkirch-Graffenstaden, France). Constitutive I-SceI expression in these cells was accomplished following transduction with MLV-ER-I-SceI-HA, which encodes a form of the endonuclease that can be translocated to the nucleus upon treatment with 4-OH-tamoxifen 3 hours post transfection (Courilleau et al., 2012). Cells were selected in medium containing puromycin at 1μg / ml and bulk cultures were expanded under selection conditions. I-SceI expression was confirmed by Western blotting of whole cell lysates using the anti-HA antibody (HA.11 clone 16B12, Covance).

